# Glial AP1 promotes early TBI recovery but chronically drives tauopathy

**DOI:** 10.1101/2021.06.17.448817

**Authors:** China N. Byrns, Janani Saikumar, Nancy M. Bonini

## Abstract

The emergence of degenerative disease after traumatic brain injury is often described as an acceleration of normal age-related processes. Whether similar molecular processes occur in injury and in age is unclear. Here we identify a functionally dynamic and lasting transcriptional response in glia, mediated by the conserved transcription factor AP1. In the early post-TBI period, glial AP1 is essential for recovery, ensuring brain integrity and animal survival. In sharp contrast, chronic AP1 activation promotes human tau pathology, tissue loss, and mortality. We show a similar process activates in healthy brains with age. Importantly, AP1 activity is present in moderate human TBI, decades after injury, and associates with microglial activation and tauopathy. Our data provide key molecular insight into glia, highlighting that the same molecular process drives dynamic and contradictory glia behavior in TBI, first acting to protect but chronically promoting disease.

## Introduction

Traumatic brain injury (TBI) has long been suggested to promote an advanced age state of the brain^1^. Both injury and age herald neurodegenerative pathology and disease, though on dramatically different timelines^2,3^. In professional athletes, disease usually manifests as chronic traumatic encephalopathy^4,5^. But select hallmarks of degeneration —tissue loss, gliosis, and aggregates of hyperphosphorylated tau protein—are also found months after a single moderate injury in otherwise healthy individuals^3,6-9^. These early changes do not guarantee clinical disease^2^, but they influence the continued trajectory of brain health. Despite the parallels between TBI and age, whether parallel cell and molecular processes occur is entirely unclear.

Glia have an emergent role in the pathogenesis of age-onset degenerative diseases^10^. As guardians of the CNS, glia maintain homeostasis by remedying pathologic and physiologic disturbances. But glia adopt aberrant states in advanced age and in disease^11,12^, with growing evidence for deleterious and degenerative impact on neurons. Alterations in genes linked to distinct neuropathologies, like *TREM2, APOE4* and *TARDBP*, commonly modify glia behavior to accelerate protein aggregation^13-16^. Inflammatory processes within glia can seed neuronal protein pathology^17,18^. Glia also propagate disease protein pathology between neurons^19,20^.

The role of glia in TBI is far less clear and often appears paradoxical^21^. In early TBI recovery, astrocytes and microglia have been reported to serve a beneficial role, though underlying intracellular processes have not been defined^21^. However, glia remain morphologically activated for decades after injury^22^. This dystrophic state, termed chronic gliosis, coincides with regions of eventual tissue loss and tau pathology^23^. Such observations hint that glia instigate degeneration long after injury, but a mechanism has yet to be identified^24^. Overall, our understanding of the molecular status of glia in TBI is incomplete, including chronic gliosis and its degenerative ramifications.

Many discoveries fundamental to human biology and disease have been made first in *Drosophila*^11,25-35^. To investigate the molecular consequences of TBI, we used a head-specific *Drosophila* TBI approach, termed dTBI^36^. Fly TBI models overcome limitations in mammals^37^ and offer exceptional genetic tractability and a short lifespan. The fly and mammalian brain have analogous core elements; namely, a repertoire of glial and neuronal subtypes, organized into functionally distinct regions and protected by a “blood”-brain barrier and a hard cuticle^38,39^. The dTBI method uses machine-controlled, tunable head compression. As such, brain degeneration progresses along a highly reproducible timeline, facilitating precise delineation of conserved molecular changes.

Here, using this highly-reproducible TBI model, we demonstrate a dynamic, lasting response in glia, mediated by the conserved transcription factor AP1. Glial AP1 has an established role in Wallerian degeneration^40,41^—a specific process to resolve axonal lesion. However, we show that AP1 activation after TBI trauma is mechanistically distinct, with implications for glia function and disease pathogenesis. In early injury, AP1 orchestrates a protective response that maintains brain integrity and ensures animal survival. In sharp contrast, sustained AP1 activity promotes pathologic hallmarks of human tau. Intriguingly, a similar process activates in glia with normal brain aging. We identify an AP1 response in humans, decades after moderate TBI, which correlates with microglial activation and tauopathy. These findings indicate that AP1 drives paradoxical behavior of glia in TBI and possibly age, first acting to protect but, in time, promoting disease progression.

## Results

### Evidence of lasting AP1 transcriptional activity after TBI

To investigate molecular drivers of degeneration after traumatic brain injury, we mapped the transcriptional time course of the dTBI response. We compared gene expression by RNAseq in sham and severe dTBI heads at 9 timepoints spanning from 1 hour post-injury (hpi) to 15 days post-injury (dpi; **Fig. 1a**). By 15 dpi, the brain is considerably degenerated^36^ and survival approximates 50% (**Extended Data Fig. 1a**). In the first 24 hpi, hundreds of genes were affected, and the number of genes up or downregulated was roughly comparable at each time point (FDR<0.05; **Fig. 1b**; **Extended Data Fig. 1b**; **Supplementary Table 5**).

**Fig. 1.**
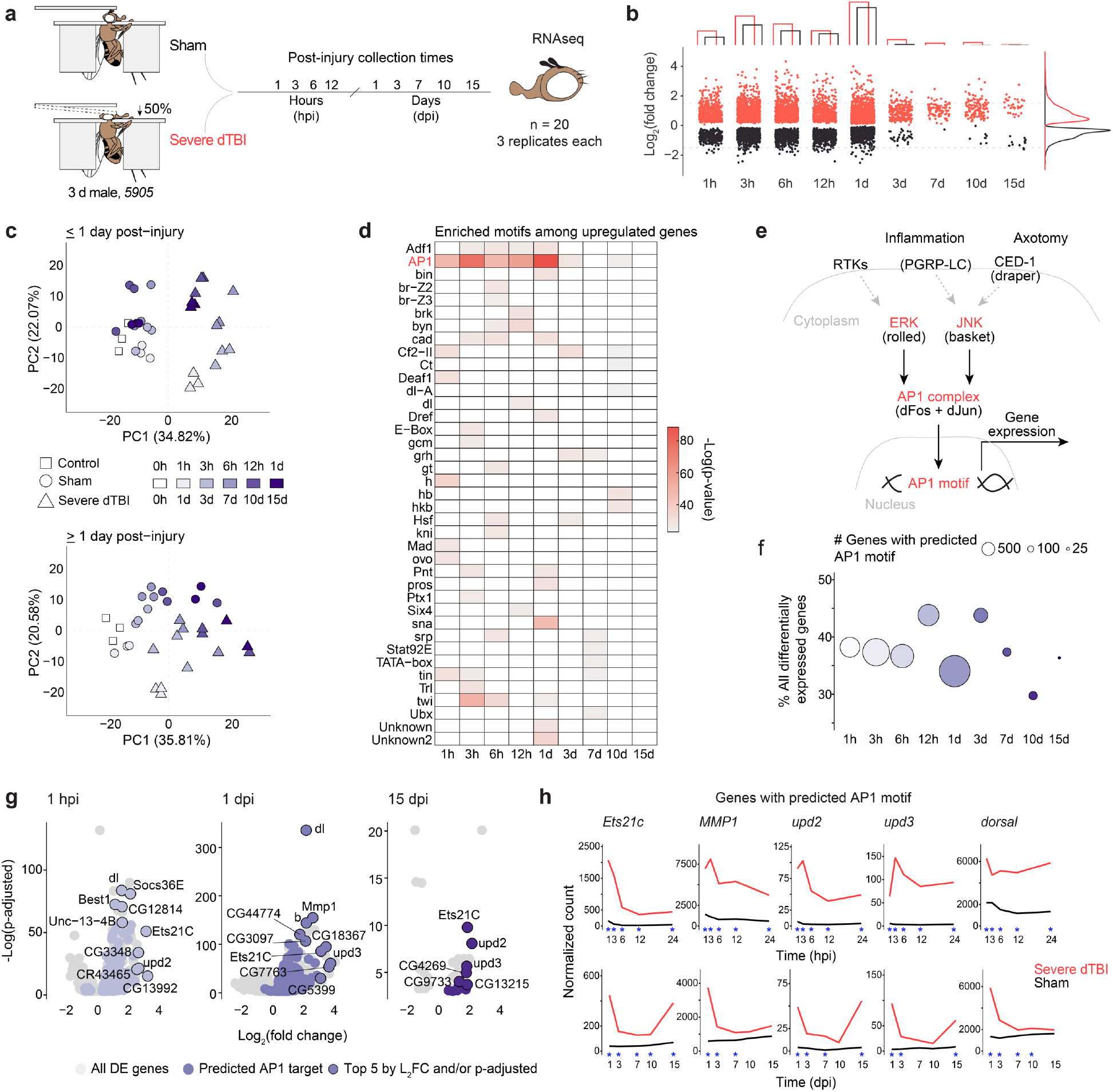
A lasting AP1 transcriptional response to TBI. **a**, Design of time course RNA-seq experiment. Sham and severe dTBI heads were collected at 1 pre-injury (0 hours per injury; hpi) and 9 post-injury timepoints as indicated with 3 biological replicates (n = 20 heads per replicate) for each injury condition and time (n = 57 samples total). **b**, P lot showing the number significantly differentially expressed genes (upregulated in red, downregulated in black) at each post-injury time. Bar graph annotation (top) summarizes the total number of genes up and down. Density plot annotation (right) summarizes the distribution by log_2_ fold change (each point = 1 gene; FDR < 0.05). **c**, Principal component analysis of ≤1 dpi (days post-injury; top) and ≥1 dpi samples (bottom). Each point represents 1 biological replicate. Shape encodes condition. Color encodes post-injury time. **d**, Tile plot showing results of HOMER *de novo* motif enrichment analysis among upregulated genes (FDR<0.05). Presence of a colored tile indicates motif enrichment at a given post-injury time. Tile color encodes significance. **e**, Regulation of the conserved transcriptional complex AP1. *Drosophila*-specific gene names in parenthesis. **f**, Summary of the proportion of all differentially expressed genes with a predicted AP1 motif. Color key as in C. **g**, Volcano plots show highlighting genes with predicted AP1 motifs (purple points), with the top 5 genes by L2FC/p-adjusted labelled. **h**, Average sham (black) and severe dTBI (red) expression of highly affected AP1 target genes at ≤1 dpi (top) and ≥1 dpi (bottom; blue star indicates FDR < 0.05 at a given time). See Supplementary Table 1 for genotypes.

By 3 dpi, the transcriptional response was substantially attenuated, and affected genes were skewed towards activation. This trend of disproportional upregulation continued through 15 dpi, with 37 of 45 genes upregulated. Notably, ∼1/3 of 15 dpi genes were also upregulated at earlier timepoints, suggesting a subset of genes may be persistently active (**Extended Data Fig. 1b**). Principal component analysis complimented and refined this characterization (**Fig. 1c**). In the first 24 hpi, dTBI and sham animals segregated across PC1 and replicates tightly clustered. After 1 dpi, samples distributed along PC1 by time, consistent with overall transcriptional resolution, though dTBI animals remained right-shifted relative to age-matched shams.

The large number of immediate gene changes, and subset of lasting gene changes, suggested key transcription factors may orchestrate the dTBI response. To uncover potential regulators, we analyzed the promoters of up and downregulated genes for *de novo* DNA motifs. A unifying motif did not emerge from downregulated genes (**Extended Data Fig. 1c**). However, upregulated genes at nearly all times were unexpectedly enriched for a common motif, predicted to be targeted by the transcriptional complex AP1 (**Fig. 1d**).

AP1 is a conserved, dimeric complex typically comprised of one member each of the Fos and Jun protein families^42^. *Drosophila* have a single member of each, referred to here as dFos and dJun (**Fig. 1e**), with structural and functional homology to mammalian Fos and Jun^43-45^. We compiled all predicted AP1 target genes to estimate their contribution to the transcriptional response. Between 30-45% of affected genes had an AP1 motif at all timepoints (**Fig. 1f**), suggesting AP1 may mediate a substantial and continued response to injury in the fly brain.

We examined the predicted AP1 targets. In the first 24 hpi, targeted genes mapped to immune and endocytic pathways (**Extended Data Fig. 1d**). This shifted to immune and JAK/STAT signaling at later times. The most highly affected genes included known fly AP1 target genes—*Ets21c, upd2, upd3, MMP1*^41,46,47^—and genes with a previously unrecognized AP1 motif, like *dorsal*, an NFkB homolog (**Fig. 1g**). Strong upregulation of these specific genes was noted throughout the post-injury period (**Fig. 1h**). By contrast, canonical stress response genes, such as the heat shock proteins and glutathione transferases, normalized to baseline by 3 dpi (**Extended Data Fig. 1e**,**f**).

### AP1 activation is a lasting, severity-dependent TBI response

To investigate AP1 activity in the brain, we used an established *in vivo* reporter line, *TREdsRed*, that expresses dsRed under the control of 4x AP1 motifs^48^. We compared changes in dsRed by mRNA and protein in dissected brains of mild and severe dTBI flies through 15 dpi; mild vs severe insults convey dramatically different mortality (**Fig. 2a**), functional impairment and brain degeneration during this period^36^. By realtime quantitative PCR (RT-qPCR), there was lasting, severity-dependent upregulation of *dsRed* (**Fig. 2b**), as well as RNA-seq predicted AP1 target genes (**Extended Data Fig. 2a**). By whole mount immunofluorescence (IF), dsRed protein increased by 1 dpi and lasted through 15 dpi (**Fig. 2c**,**d**). dTBI did not affect dsRed with a mutated AP1 motif, *MREdsRed*^48^, confirming AP1 specificity (**Extended Data Fig. 2b**). Thus, AP1 activity appears to be an enduring consequence of dTBI. This occurs even after mild injury, where injured flies are indistinguishable from shams in behavior, but develop degeneration with time^36^.

**Fig. 2.**
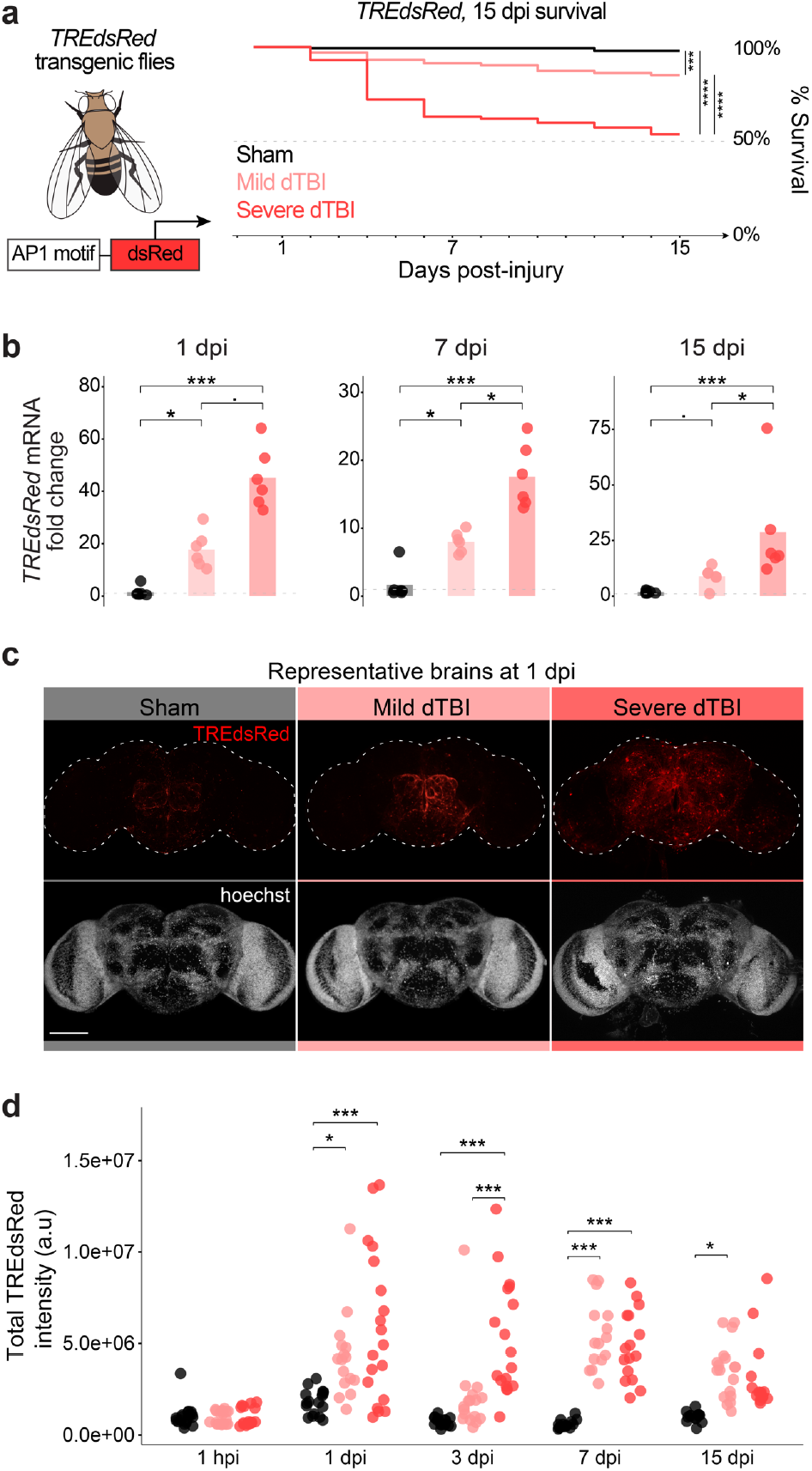
AP1 activation is sustained and severity-dependent. **a**, AP1 activation was assessed in *TREdsRed* transgenic flies, with dsRed under 4X AP1 motifs. Right, 15 dpi survival of *TREdsRed* flies injured with sham (black), mild (35% head compression), or severe (45% head compression) dTBI (n = 100 per condition, 5 vials of 20 flies; p < 0.0001, Kaplan-Meier analysis with log-rank comparison). **b**, Mean relative expression of *dsRed* mRNA by RT-qPCR at 1, 7, and 15 dpi between injury conditions (each point = 1 biological replicate of 9 dissected brains; n = 6 biological replicates per condition; Kruskal– Wallis test with Dunn’s multiple comparison test and Holm adjustment). See Extended Data Fig. 2a for additional AP1 target genes. **c**, Representative z-stacked whole mount brains at 1 dpi across injury conditions. Scale bar represents 100 *μ*m. **d**, Quantification of dsRed immunofluorescence in whole mount brains throughout the post-injury period (each point = 1 brain; n = 14-19 brains per condition/time pooled from two independent experiments; p = 3.33e-09, two-way ANOVA with Tukey’s test). Statistical annotations are ****p<0.0001, ***p<0.001, **p<0.01, *p<0.05. All scale bars 100*μ*m.

### AP1 activates and persists in glia

To localize AP1 activation within the brain, we examined *TREdsRed* whole mount brains by high magnification confocal microscopy. Surprisingly, the pattern of dsRed strongly resembled glial processes and not neurons (**Fig. 3a**), the latter of which constitute most cells (∼90%) in the fly brain^38^. We therefore quantified dsRed in glia (repo+) and neurons (elav+) by whole mount IF after mild and severe dTBI.

**Fig. 3.**
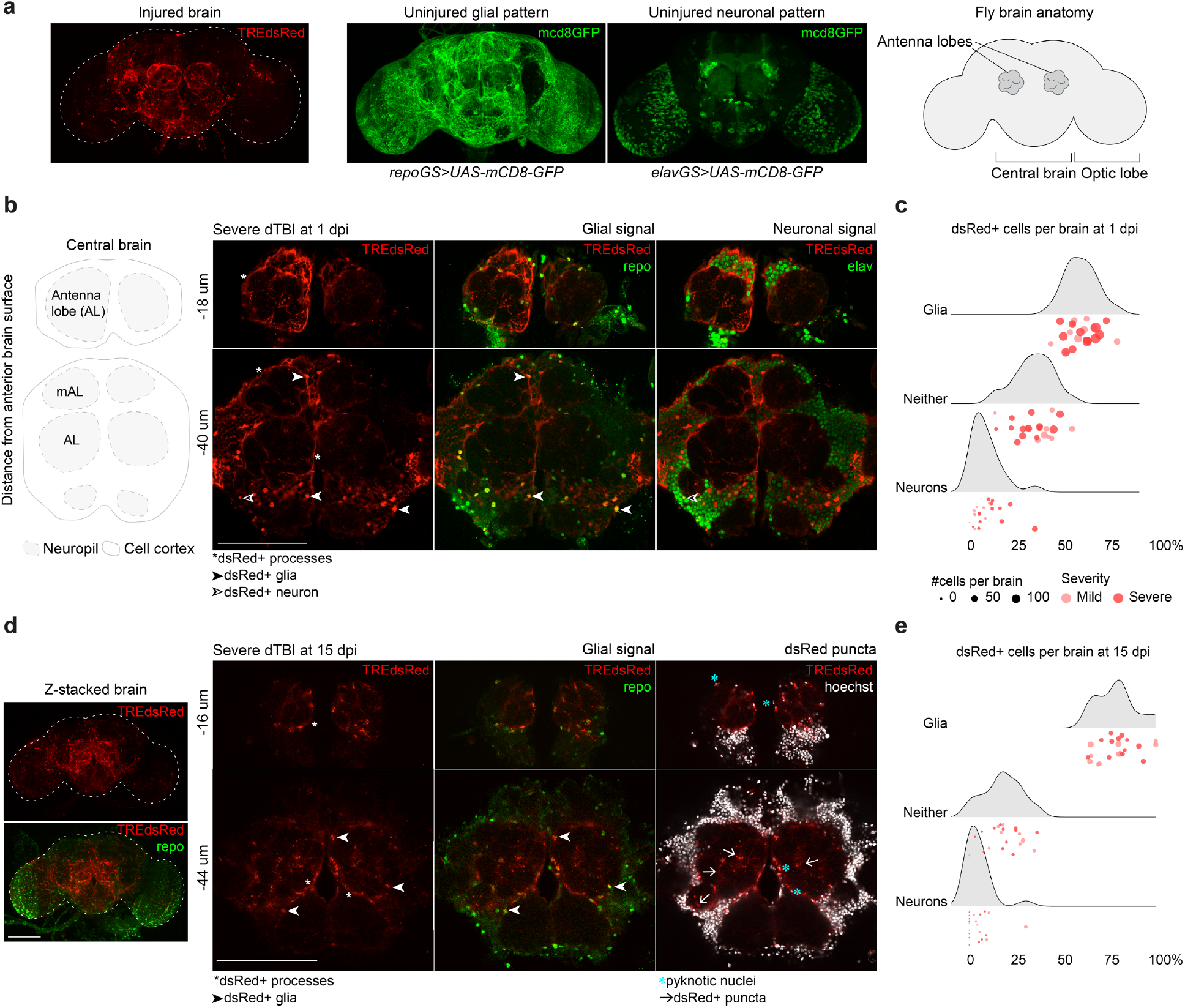
AP1 activates and persists in glia. **a**, Representative z-stacked whole mount brain showing dsRed pattern in an injured brain (severe dTBI, 1 dpi). Also shown are the pattern of glia and neuron membranes in uninjured brains, highlighted using a membrane-targeted GFP variant (mcd8-GFP) expressed under an inducible glia (*repoGS* aka *repoGeneSwitch*) or neuron-specific (*elavGS* aka *elavGeneSwitch*) promoter. Simplified anatomy of the fly brain on the right, as seen in whole mount brain images. **b**, Representative high-magnification images of the central brain at 1 dpi. Thick, hypertrophic dsRed+ membrane processes are detected throughout the brain. Many dsRed+ cells co-localize with a nuclear glia-specific antibody (repo). **c**, Ridgeline plot demonstrating the proportion of dsRed+ cells that are glia (repo+), neurons (elav+) or neither (repo-/elav-) in mild (pink) and severe (red) dTBI brains (each point = 1 brain). X-axis indicates the percentage of all dsRed+ cells. Y-axis indicates sample density along the x-axis. **d**, Representative high-magnification images of the central brain at 15 dpi. dsRed+ processes are thinner and appear fragmented (see white arrow heads). dsRed+ repo+ nuclei appear shrunken (see blue *). Anucleated dsRed+ puncta are abundant throughout the neuropil (see white arrows). On the left is the whole z-stacked brain, demonstrating the overall pattern of dsRed in late dTBI. **e**, Ridgeline plot as in C for 15 dpi dTBI brains. All scale bars 100*μ*m. See Supplementary Table 1 for genotypes.

At 1 dpi, most dsRed+ cells were repo+ glia (59.99 ± 8.51%) and only a minority of cells were elav+ neurons (8.99 ± 8.01%; **Fig. 3b**,**c**). Severe dTBI brains on average had more dsRed+ cells than mild dTBI brains (159 vs 83), though the proportion of dsRed+ glia to neurons was comparable. Overall, AP1 activity was greatly enriched in glia, representing ∼6-fold greater proportion of cells compared to neurons (Fisher’s exact test, p = 2.53e-33). dsRed+ glia appeared morphologically reactive, with characteristic thick, hypertrophic processes and were concentrated around primary impact areas, like the antenna lobes (see **Fig. 3b**).

By 15 dpi, the total number of dsRed+ cells per brain decreased (from 125±53 to 31±12), though a greater percentage were repo+ glia (78.77±10.70%; **Fig. 3e**). Key histological differences further distinguished dsRed+ repo+ glia at 15 dpi from 1 dpi (**Fig. 3d**). dsRed+ glia had pyknotic nuclei and irregular processes. Anucleated dsRed+ puncta were present throughout the neuropil and around repo+ nuclei, suggestive of cytoplasmic fragmentation. Altogether, these changes are reminiscent of dystrophic gliosis, a morphological phenomenon of chronically active mammalian glia^49,50^ previously unreported in flies^38^. These morphological hallmarks appeared as early as 3 dpi after mild and severe dTBI but were predominant by 7 dpi. Thus, a fraction of glia transition to an altered state, defined by their morphology and chronic activation of AP1.

### Activation of glial AP1 in TBI is distinct from Wallerian degeneration

Glial AP1 is integral to Wallerian degeneration, an active degeneration process following axonal lesion^40,51^. In flies, glia sense this neuron-specific injury via the CED1 receptor, draper, without which glia fail to upregulate AP1 target genes and fail to clear axonal debris^40,41^. We investigated if the AP1 response in dTBI is draper-dependent, by eliminating *draper* gene function in glia and assessing AP1 activity via TREdsRed. Unlike antennal nerve lesion (AL) injury^51^, where *draper-RNAi* prevented dsRed expression and glial hypertrophy (**Extended Data Fig. 3a**), *draper-RNAi* had no effect on dsRed protein or mRNA in dTBI (**Extended Data Fig. 3b**,**c**). As draper detects the molecular signature of axonal debris, glia seemingly answer to distinct cues with dTBI.

Two mechanisms are known to promote the transcriptional activity of AP1. Dimer stabilization via phosphorylation by MAP kinases (MAPK), and increased levels of component proteins^52,53^. In Wallerian degeneration, AP1 is phosphorylated by the c-Jun terminal kinase (JNK/basket; see **Fig. 1e**), which acts downstream of draper^40^. To determine if JNK also regulates AP1 in dTBI, we assessed active pJNK, JNK, and dsRed by western immunoblot in *TREdsRed* brains at 30 min post-injury (mpi), 1 hpi, 1 dpi, and 15 dpi. Though dsRed increased, we failed to detect a concomitant change in pJNK/JNK (**Fig. 4a**; **Extended Data Fig. 3d**). Genetic reduction of JNK with *basket* RNAi failed to mitigate dTBI-induced upregulation of dsRed (**Fig. 4b; Extended Data Fig. 3e)**, despite validation of RNAi efficiency to reduce *dsRed* and *basket* mRNA in whole flies under a ubiquitous driver (**Extended Data Fig. 3f**). Thus, unlike Wallerian degeneration, glial AP1 activation in TBI appears to be both draper and JNK-independent.

**Fig. 4.**
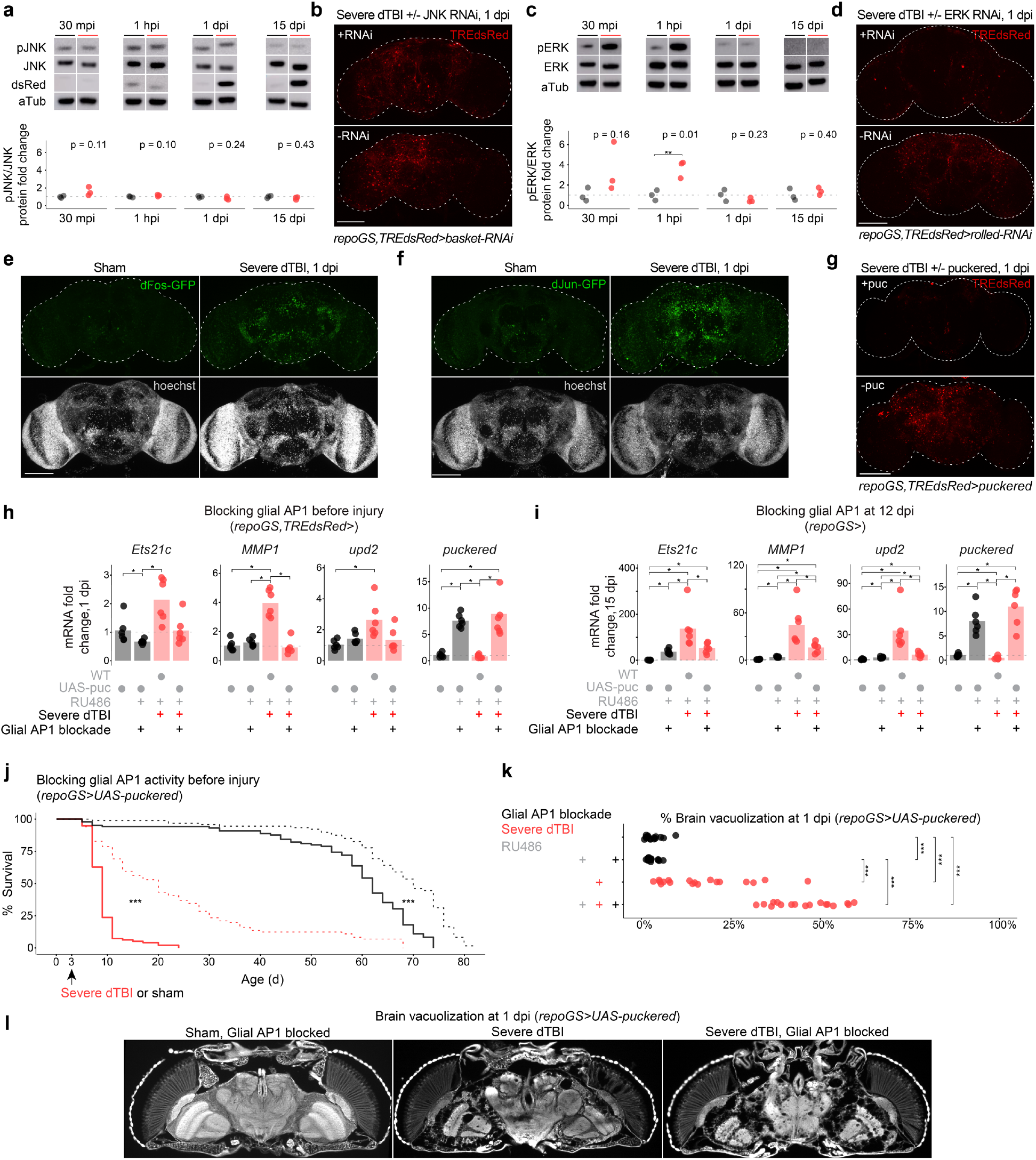
Glial AP1 is activated by ERK and is necessary for early TBI recovery. **a**, Representative protein changes (top) and quantification of pJNK/JNK ratio by western immunoblot (each point = 1 biological replicate, 8 dissected brains; n = 3 biological replicates per condition; Student’s t-test for each timepoint). See Extended Data Fig. 3d for immunoblots of quantification. **b**, Endogenous dsRed protein in whole mount brains at 1 dpi, with (top) or without (bottom) RNAi against JNK expressed under a glia-specific inducible promoter. See Extended Data Fig. 3e for quantification. **c**, Representative protein changes (top) and quantification (each point = 1 biological replicate, 8 dissected brains; n = 3 biological replicates per condition; Student’s t-test for each timepoint). See Extended Data Fig. 4b for immunoblots of quantification. **d**, Endogenous dsRed protein in whole mount brains at 1 dpi, with (top) or without (bottom) RNAi against ERK expressed under a glia-specific inducible promoter (representative of n = 7-9 brains each). **e**, Representative whole mount brains at 1 dpi in flies expressing dFos-GFP under the endogenous dFos promoter. Representative of n = 9 – 16 brains per condition. **f**, Representative whole mount brains at 1 dpi in flies expressing dFos-GFP under the endogenous dJun promoter. Representative of n = 8 – 13 brains per condition. **g**, Representative whole mount brains at 1 dpi, with (top) or without (bottom) puckered expressed under a glia-specific inducible promoter. See Extended Data Fig. 4c for quantification. **h**, Mean relative expression of AP1 target genes by RT-qPCR at 1 dpi, with or without AP1 activity blocked in glia via *puckered* expression (each point = 1 biological replicate, 9 dissected brains; n = 6 biological replicates per condition; Kruskal–Wallis test with Dunn’s multiple comparison test and Holm adjustment). **i**, Mean relative expression of AP1 target genes by RT-qPCR at 15 dpi, with or without AP1 activity blocked in glia beginning at 12 dpi via *puckered* expression (each point = 1 biological replicate, 9 dissected brains; n = 6 biological replicates per condition; Kruskal–Wallis test with Dunn’s multiple comparison test and Holm adjustment). **j**, Post-injury survival with (RU; dashed line) or without (vehicle; solid line) AP1 activity blocked in glia via *puckered* expression (n = 100 per condition, 5 vials of 20 flies; p < 0.0001, Kaplan-Meier analysis with log-rank comparison). **k**, Representative brain vacuolization at 1 dpi under sham and dTBI conditions, with or without AP1 activity blocked in glia via *puckered* expression. Quantification on right, expressed as % of total brain area that is vacuolized (each point = 1 brain; n = 15-18 brains per condition pooled from two independent experiments; p = 5.07e-10, two-way ANOVA with Tukey’s test). Statistical annotations are ****p<0.0001, ***p<0.001, **p<0.01, *p<0.05. All scale bars 100*μ*m.

### ERK is necessary for glial AP1 activity in TBI

To identify the upstream activator of AP1, we considered the MAPK extracellular signal-regulated kinase, ERK. ERK phosphorylates dFos and dJun at sites distinct from JNK^54,55^. *Rolled*, the gene encoding fly ERK, is also detected in a greater a proportion of glia than *basket* at baseline (**Extended Data Fig. 4a**). Active pERK and ERK were assessed by western immunoblot after dTBI. pERK/ERK ratios mildly increased at 30 mpi, strongly increased at 1 hpi then normalized by 1 dpi (**Fig. 4c; Extended Data Fig. 4b**). RNAi knockdown of *rolled* in glia entirely eliminated cellular dsRed signal (**Fig. 4d**). Altogether, these findings indicate ERK is activated by dTBI and is required for activation of glial AP1.

AP1 is a dimer, typically comprised of Fos and Jun. By RNAseq, *dFos* and *dJun* expression increased with injury (**Extended Data Fig. 4c**). GFP-tagged variants of dFos and dJun robustly increased at 1 dpi (**Fig. 4e**,**f**), indicating that the protein levels are also increased and suggesting that AP1 is a dFos/dJun heterodimer. To assess the function of these AP1 components, we examined the effects of *puckered*, a MAPK phosphatase shown to reduce dFos and dJun protein^56^. As with ERK RNAi, *puckered* expression in glia eliminated cellular dsRed signal (**Fig. 4g; Extended Data Fig. 4c**). *Puckered* did not alter pERK levels, consistent with an effect on AP1 and not ERK activity (**Extended Data Fig. 4d**). Expression of *puckered* in glia also suppressed dTBI-induced upregulation of AP1 target genes by RT-qPCR at 1 dpi (**Fig. 4g**). These data indicate that perturbation of AP1 in glia can abolish select dTBI-induced gene changes. To determine if AP1 continues to drive gene expression in late injury, we expressed puckered at 12 dpi then assessed AP1 target genes by RT-qPCR at 15 dpi. Gene levels were reduced by more than 50% in dTBI though expression was not restored to sham levels (**Fig. 4I**). Thus, AP1 appears to drive chronic gene expression in glia.

### Glial AP1 promotes early TBI recovery

The precise role of glia in TBI, whether helpful or harmful, is unclear and may depend on both context and duration of glia activation^21^. To define the function of AP1 in the early dTBI period (<3 dpi), when glia were strongly hypertrophied and without dsRed+ puncta, we blocked glial AP1 activity via *puckered* before injury and assessed animal survival. Mean and maximum lifespan were dramatically reduced in dTBI animals absent of glial AP1 activity (**Fig. 4j**). Curiously, shams were also affected, but only late in age. To distinguish mortality due to a loss of AP1 target genes from other possible effects of *puckered*, we blocked the AP1 target *Ets21c*, which was highly expressed in glia after dTBI (**Extended Data Fig. 5a**). RNAi knockdown of *Ets21c* in glia phenocopied mortality, resulting in hastened death after dTBI (**Extended Data Fig. 5b**). Altogether, these findings indicate dTBI would be rapidly lethal in the absence of glial AP1 activation.

Neurodegeneration, which manifests as vacuolization in the fly brain, progresses gradually after dTBI and in age^36^. However, blocking glial AP1 with puckered resulted in extraordinary degeneration by 1 dpi, with ∼50% loss of brain tissue (**Fig. 4k**,**l**). Vacuoles were concentrated in cortical layers rather than the neuropil, suggestive of loss of neuronal cell bodies. Large, coalescent vacuoles formed as early as 30 mpi (not shown), indicating that preventing glial AP1 activation has a remarkable effect to promote neuronal loss. We next examined the effects of *puckered* expression in neurons, given that some neurons appeared dsRed+ after dTBI. Neither brain vacuolization nor animal survival (**Extended Data Fig. 5c**,**d**) were appreciably altered, underscoring a highly glia-specific role for AP1. Emphasizing the ERK-specificity of glial AP1 activity in dTBI, targeting glial JNK also had no effect on brain vacuolization (**Extended Data Fig. 5e**). Altogether, these findings suggest glial AP1 is highly protective in early dTBI and without it, brain injury is catastrophic.

### Glial AP1 activates with age in healthy brains

As noted, TBI is hypothesized to accelerate normal brain aging processes^57,58^: survivors develop cognitive impairment and pathology associated with advanced age and are at risk for age-onset degenerative disease^2,6,59,60^. But whether common molecular processes govern the seemingly analogous changes in TBI and in age is unclear.

We observed in the PCA analysis of late post-injury RNAseq data (see **Fig. 1c**) that dTBI survivors were right-shifted compared to shams, suggestive of an “older” transcriptome. To investigate this, we compared genes upregulated in the fly brain with age^61^ to those upregulated in late dTBI (**Fig. 5a**). Age-associated genes showed a 6.4-fold enrichment in late injury (Fisher’s exact test, p = 2.73e-17), consistent with dTBI brains showing accelerated upregulation of age-associated genes. Even more striking, 40% of shared genes were predicted AP1 targets (n = 13/32). To determine if AP1 genes are upregulated in age, we performed RT-qPCR for common AP1 targets in 5 vs 40 d brains of reporter *TREdsRed* flies. Intriguingly, all assayed genes showed a robust increase with age (mean fold change 2.3-7.4; **Fig. 5b**). Consistent with an impact on gene expression, dsRed protein in 40 d brains was abundant by whole mount IF and, as with dTBI, largely localized to glia (**Fig. 5c**). Remarkably, aged dsRed+ glia also morphologically resembled dsRed+ glia in late dTBI, with noticeable dsRed+ puncta (**Fig. 5d**). Altogether, these data suggest age and TBI evoke a similar response by glia, marked by dystrophy and AP1 activation.

**Fig. 5.**
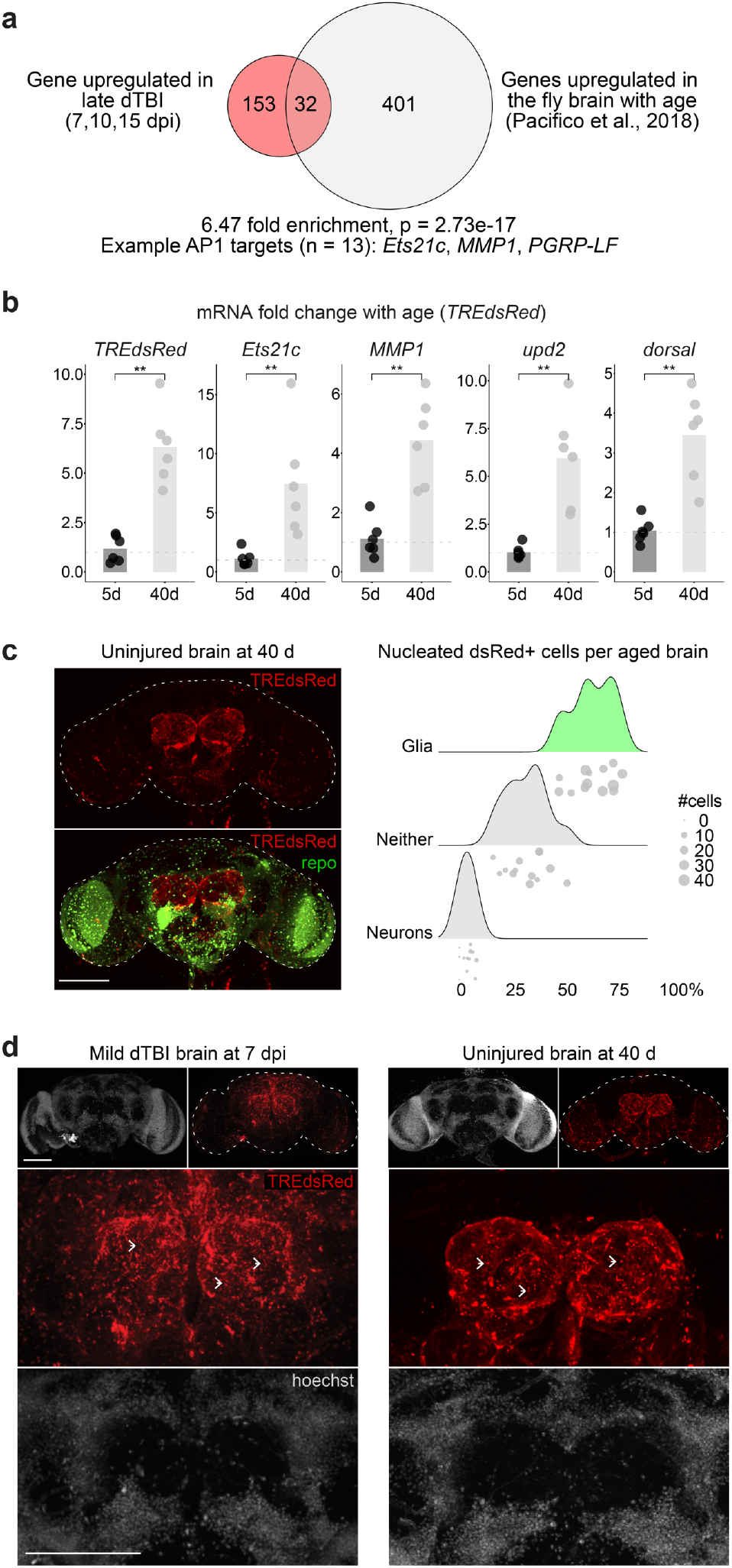
Age and injury evoke a glial AP1 response. **a**, Enrichment of genes upregulated in late dTBI and with age (0 vs 40 d male) in the fly brain (one-sided Fisher’s exact test). 40% of overlapping genes are predicted AP1 target genes (n = 13). **b**, Mean relative expression of select AP1 target genes mRNA by RT-qPCR in 5 vs 40 d old brains (each point = 1 biological replicate, 9 dissected brains; n = 6 biological replicates per condition; Wilcoxon test for each gene). **c**, Left, representative z-stacked whole mount brain of uninjured 40 d old brain. Right, ridgeline plot, as shown in Fig.s 3C and 3E, showing the proportion of dsRed+ cells that are glia (repo+), neurons (elav+) or neither in 40 d old brains (each point = 1 brain). **d**, Representative z-stacked whole mount brains: left, 7 dpi brain, right, 40 d old uninjured brain. Not shown is dsRed signal in 5 d uninjured brains, which is identical to shams at 1 dpi, see Fig. 2b. White arrows highlight anucleated dsRed+ puncta. Statistical annotations are ****p<0.0001, ***p<0.001, **p<0.01, *p<0.05. All scale bars 100*μ*m. See Supplementary Table 1 for genotypes.

### Chronic glial AP1 activation promotes human tau pathology

Pathological aggregates of hyperphosphorylated tau protein emerge with age and in age-onset degenerative diseases; they appear within months following moderate TBI^4,6,7,22,62^. Notably, human tau forms pathologic aggregates in flies with age^63^. In TBI patients and animal models, tau pathology colocalizes with chronically activated microglia and astrocytes^6,7,22,23,64-67^. Thus, we considered that sustained glial AP1 activity after dTBI may promote human tau pathology.

We expressed human wild-type tau (0N4R isoform) in glia and subjected animals to injury. Paraffin embedded heads were examined for phosphorylated tau over time. While 1 dpi showed no immunoreactivity, by 5 dpi, dTBI brains showed dramatic puncta positive for AT100 or AT8, two distinct pathological phosphorylation sites on tau (**Fig. 6a**). Phospho-tau puncta did not form after injury when tau was expressed in neurons (**Extended Data Fig. 6a**). Increased puncta upon glial expression coincided with a size shift of tau to a higher molecular weight by semi-denaturing gel electrophoresis; this resolved with phosphatase treatment (**Extended Data Fig. 6b**) indicating the formation of phosphorylated tau oligomers. There was also a positive correlation between puncta and brain vacuole area (**Extended Data Fig. 6c**), suggesting tauopathy exacerbates tissue loss. Puncta appeared concentrated around multi-vesicular vacuoles in the neuropil (**Extended Data Fig. 6d**), similar to dsRed+ puncta. The functional impact of tauopathy was assessed in mild injury, as tau was highly toxic with severe dTBI with most animals expiring soon after 10 dpi. Mild dTBI mortality was significantly increased, indicating tau pathology is injurious and exacerbates dTBI (**Fig. 6b**).

**Fig. 6.**
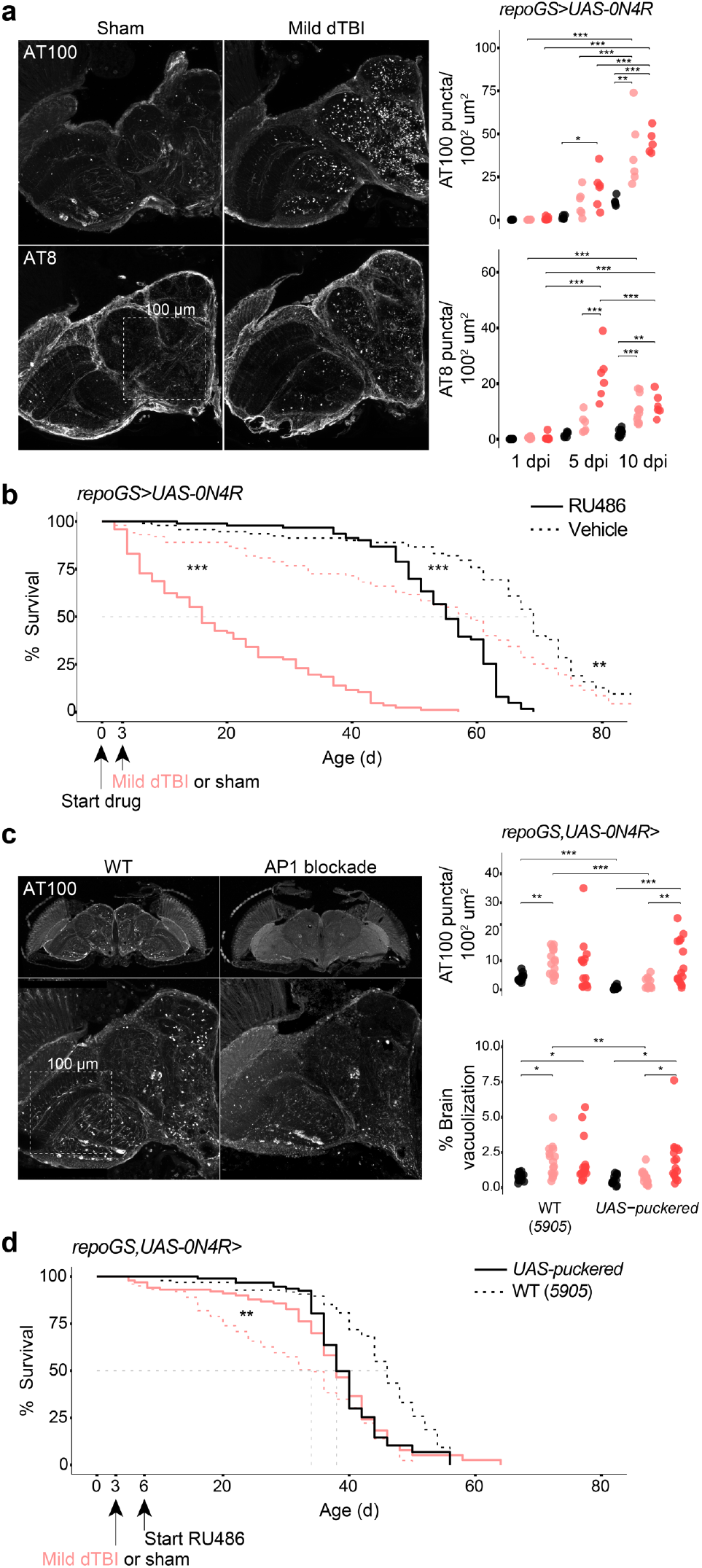
Chronic AP1 activity promotes human tau pathology. **a**, Left, representative hemisection of paraffin-embedded heads at 10 dpi, immunostained for tau phosphorylation sites, AT100 and AT8. Box illustrates area of 100^2^ *μ* m^2^. Right, quantification of the number of AT00+ and AT8+ puncta, represented as puncta per 100 *μ*m^2^ at 1, 5 and 10 dpi (each point = 1 brain, n = 6 per condition/time pooled from two independent experiments; AT8: p = 6.40e-05, AT100: p=0.000336, two-way ANOVA with Tukey’s test). **b**, Post-injury survival with (RU; solid) or without (vehicle; dashed) human tau expression in glia (n = 100 per condition, 5 vials of 20 flies; p < 0.0001, Kaplan-Meier analysis with log-rank comparison). **c**, Left, representative z-stacked hemisections of paraffin-embedded sham and mild dTBI heads at 10 dpi, immunostained for AT100. Right, quantification of AT00+ as in 6A (top) and % brain vacuolization (bottom) at 10 dpi without *puckered* (WT) or with *puckered* expression in glia from 3 dpi (each point = 1 brain, n = 12-16 per condition/genotype pooled from two independent experiments; AT100: p=0.0093, brain vacuolization: p=0.068, two-way ANOVA with Tukey’s test). **d**, Post-injury survival with *puckered* (solid) or without (dashed) *puckered* expression in glia in the setting of tau expression (n = 100 per condition, 5 vials of 20 flies; p < 0.0001, Kaplan-Meier analysis with log-rank comparison). Statistical annotations are ****p<0.0001, ***p<0.001, **p<0.01, *p<0.05. All scale bars 100*μ*m. See Supplementary Table 1 for genotypes.

To determine whether glial AP1 activation modulated tauopathy, we perturbed AP1 with *puckered*. We initiated *puckered* expression at 3 dpi, when glia begin to exhibit a morphological shift to dystrophy, then quantified AT100 puncta at 10 dpi. Compared to flies expressing tau alone, *puckered* significantly reduced puncta after mild, but not severe, dTBI (**Fig. 6c**). Brain vacuolization in mild dTBI brains was also restored to sham levels (**Fig. 6c**). Blocking glial AP1 partially rescued tauopathy-exacerbated mortality in mild dTBI (**Fig. 6d**; mean survival, 31 d vs 37 d; median survival, 34 vs 38 d). After this period, mortality increased in both sham and dTBI flies expressing puckered, consistent with earlier findings (see shams in **Fig. 4j**) that targeting glial AP1 activity is deleterious in aged animals. These data indicate that blocking AP1 activity during an optimal window after mild dTBI can prevent tauopathy, along with associated degenerative and functional consequences.

Altogether, these data suggest a model (**Fig. 7**) in which trauma initiates rapid, ERK-involved AP1 activation in glia. AP1 then orchestrates a molecular response essential for early brain recovery and continued animal survival. But continued activation of glial AP1 can promote tauopathy to cause brain tissue loss and mortality

**Fig. 7.**
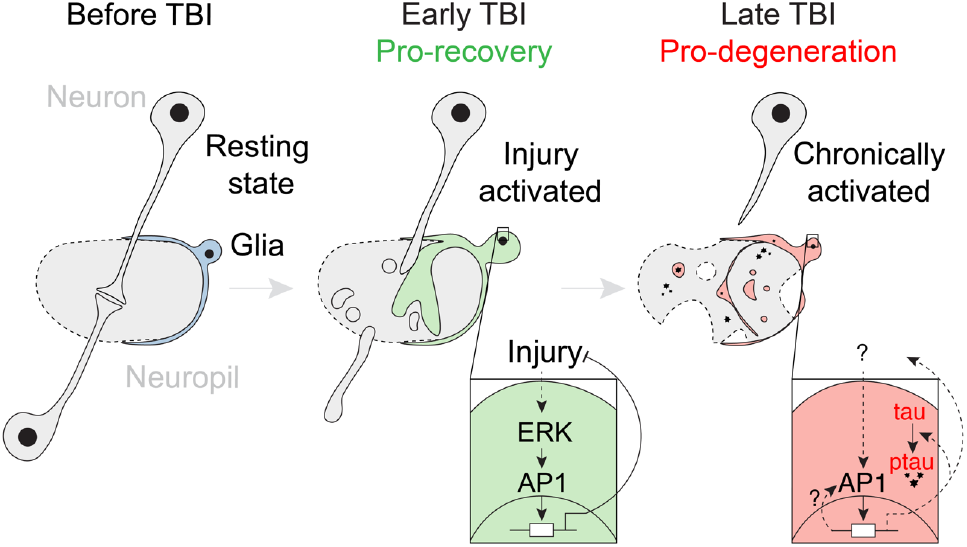
Glial AP1 promotes early TBI recovery but chronically drives tauopathy. Left, simplified fly brain schematic, showing an intracortical synapse surrounded by a resting state glia. Middle, in the early post-injury period (<3 dpi), ERK activates glial AP1, which orchestrates a gene program vital to injury recovery. Right, sustained AP1 activity fosters a prodegenerative glial state that accelerates human tau pathology. Solid lines are known interactions; dashed lines are proposed interactions.

### Evidence of AP1 activity decades after moderate TBI in humans

To extend our findings to humans, we leveraged RNA-seq and pathology data from the Allen Institute’s Aging, Dementia and TBI study. In contrast to studies that focus on CTE-confirmed athletes, donors in this cohort had a moderate injury exposure. The vast majority of injuries occurred before age 30, and unconsciousness typically lasted for less than ten minutes (**Extended Data Fig. 7a**). All donors were of advanced age (70-100+ years old) and expired from non-TBI causes.

Differential expression analysis between TBI (n = 80 samples) and non-TBI donors (n = 81 samples) showed 478 affected genes (FDR < 0.10; **Supplementary Table 6**), with more down (n = 359) than upregulated genes (n = 119). Characterization of upregulated genes using the Molecular Signature Data Base’s (MSigDB) Hallmark gene sets revealed the most enriched gene set as “TNF*α* signaling via NFkB”. The top 10 enriched gene ontology terms (p < 0.00001; **Extended Data Fig. 7c**) were dominated by processes relating to MAPK activation and protein modification, hinting at active kinase signaling with altered protein processing.

Motif enrichment analysis among upregulated genes uncovered a total of 12 motifs, two of which were AP1 component protein motifs—FRA1 (fos-related antigen 1) and JUND (**Fig. 8a**). Predicted FRA1/JUND gene targets included *JUNB, JUND, MMP9*, and *GADD45a*, as well as members of the small heat shock protein family (**Fig. 8b; Extended Data Fig. 7d**). The upregulation of JUND and JUNB genes suggests a potential feed-forward mechanism sustained AP1 activation in humans. Notably, an NFkB (*RELB*) was also targeted, as in dTBI.

**Fig. 8.**
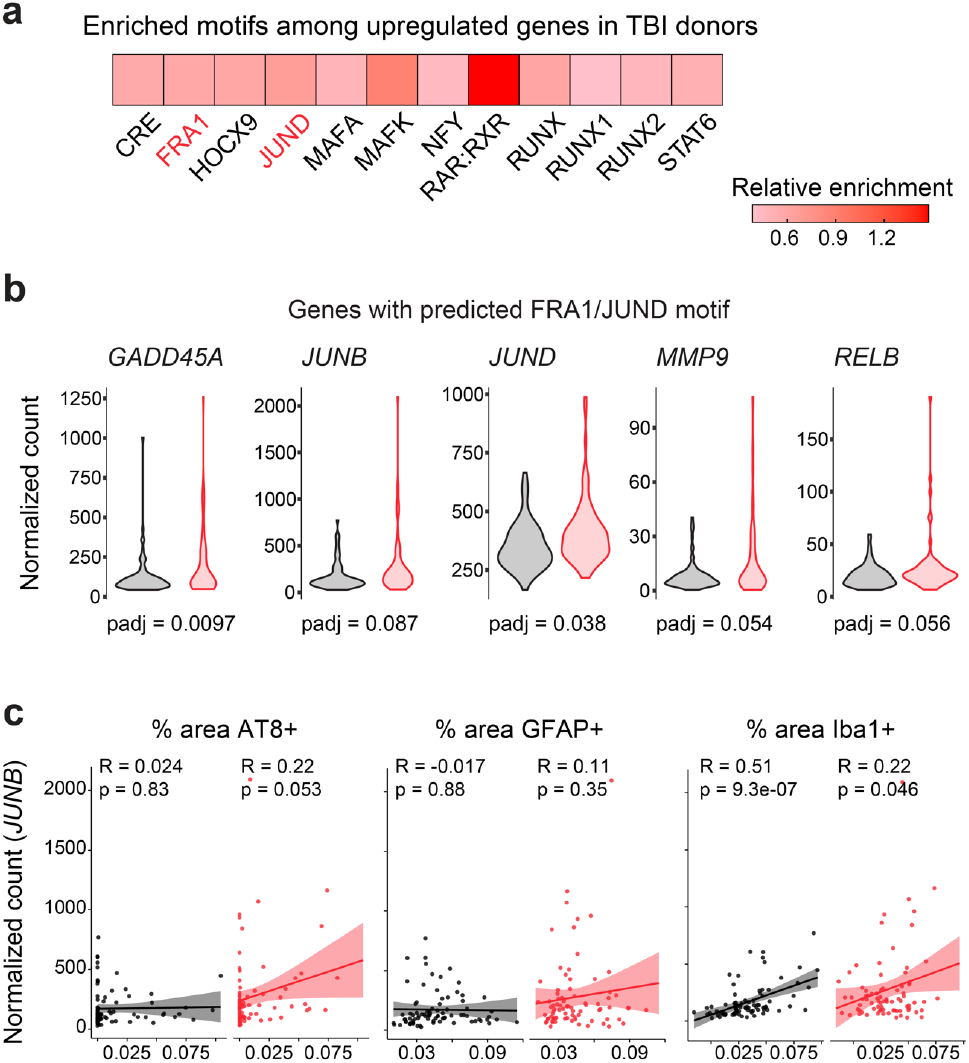
AP1 activity decades after moderate TBI in humans. **a**, Enriched motifs detected by HOMER among upregulated genes in TBI donors (FDR<0.05), colored by relative enrichment to non-TBI donors. **b**, Violin plots showing expression of select predicted FRA1/JUND genes in non-TBI (black; n = 81 samples) and TBI (red; n = 80 samples) donors (FDR<0.10). See Fig. S8D for all target genes. **c**, Pearson’s correlation between *JUNB* expression and % area of AT8, GFAP, and Iba1 by IHC in non-TBI (black; n = 81 samples) and TBI (red; n = 80 samples) donors (red; each point = 1 sample).

Among the FRA1/JUND targets, *JUNB* was among the most highly expressed genes. We examined *JUNB* to determine how transcriptional activity of FRA1/JUND may relate to pathology. Pearson’s correlation between *JUNB* expression and AT8 abundance by IHC revealed a near significant positive association in TBI donors (R = 0.22, p = 0.053; **Fig. 8c**). To assess correlation with glial activation, we examined % area by IHC positive for GFAP and Iba1, the respective markers for activated astrocytes and microglia. Regardless of injury status, *JUNB* expression positively correlated with Iba1 (**Fig. 8c**). These findings indicate that transcriptional activity of AP1 is evident decades after moderate human TBI, and positively correlates with markers of microglial activation and tau pathology.

## Discussion

We find that TBI triggers functionally dynamic and sustained glial activity, mediated by the transcriptional complex AP1. In the early post-injury period, our data suggest AP1 coordinates a protective molecular response that accompanies a hypertrophic morphological reaction, akin to glial hypertrophy described in mammalian injury^21^. Preventing AP1 activation dramatically worsens dTBI outcomes, suggesting AP1 acts to counter harmful early injury processes. These findings offer molecular insight into how glia resolve the early response to TBI, which has altogether eluded mammalian TBI studies due a lack of glia-specific genetic tools^21^.

In dTBI, we find that glial AP1 activation is draper and JNK-independent. This indicates that the molecular pathophysiology of TBI (this work) and Wallerian degeneration^40,51,68^ are distinct. In the latter case, glia sense neuronal injury indirectly via the CED-1 family of receptors. By contrast, TBI is a global trauma that directly damages neurons and glia. Downstream of draper/CED-1 in Wallerian degeneration, AP1 is activated by JNK^40,68^. Yet in dTBI, our findings support AP1 activation by ERK. These data are consistent with distinct roles for JNK and ERK, where site-specific phosphorylation of AP1 achieves distinct biological outcomes^54,55^. TBI and Wallerian degeneration likely converge downstream of AP1, as dTBI and axonal lesion injury activate similar target genes^41^. Critically, axonal lesion does not lead to chronic gliosis or chronic AP1 activation^51^, as draper-dependent processes terminate glia activity^69^. These data raise several paths by which TBI may uniquely trigger chronic glial activation.

In sharp contrast to the beneficial role of AP1 in early injury, we find that sustained AP1 activity coincides with a harmful glial phenotype, marked by dystrophic morphology and human tau pathology. Eliminating continued glial AP1 reduces tau pathology and improves brain degeneration. These findings clarify work showing that the elimination of chronically activated microglia in murine TBI is beneficial^70^. ERK was recently implicated in promoting the accumulation of c/EBP*β* in microglia, accelerating tauopathy in mice^71^. It was suggested that c/EBP*β* may interact with AP1 to target degenerative genes, representing an important overlap with the process we identify. Altogether, these data support an emerging role for an ERK-AP1 pathway in prodegenerative glial behavior. Critical next steps involve further delineation of upstream and downstream processes.

We demonstrate a positive association between TBI, FRA1/JUND activity and pathologic AT8 levels in humans. Mammalian FRA (fos-related antigen) proteins are stress-induced splice variants of *fosB*^72^, are preferentially targeted by ERK^73^, and form chronically active AP1 dimers^74^. Their activity is most robust with repeated stress^74^, with implications for repetitive TBI. Though *Drosophila* lack FRA proteins^45^, our data suggest dFos behaves like mammalian FRA in dTBI, by displaying chronic activation. To this end, the RNAseq data uncover a dTBI-specific *dFos* isoform (Byrns and Bonini, unpublished). By probing human data sets, we find AP1 activity correlates with markers of microglial activation, regardless of injury. As donors were of advanced age, this may speak to a dual role for glial AP1 in injury and age.

Our data indicate that a pro-degenerative glial phenotype emerges when AP1 activity outlasts the harmful processes they first served to counter. Along this reasoning, AP1 should be dispensable after the early injury period. Consistent with this, we find that MAPK phosphatase expression from 3 dpi and later does not impact continued dTBI survival, despite lethality when expressed prior to injury. Chronic AP1 activity may be the remnant of an early response that fails to terminate, as with positive feedback loops^75^ or highly stable transcripts^76^, and/or a consequence of continued ERK signaling.

TBI has been suggested to promote an advanced age state of the brain^60,77^. Our data indicate a molecular similarity between TBI and aging may be tied to glial AP1. Increasing evidence shows age and disease evoke abnormal glial states^12,78^. Our work adds molecular insight, identifying AP1-related transcriptional and morphological changes in glia of young injured brains and old uninjured brains. Glial AP1 perturbation hastens mortality in aged flies (shown in **Fig. 4j, 6d**), as in early dTBI. With time, AP1 may activate to counter harmful age-associated processes but remain chronically active. Other CNS insults, like extracellular amyloid accumulation, may also trigger glial AP1 activity and eventual tau pathology. These findings have critical implications for therapeutic efforts. An optimal strategy to mitigate degeneration after TBI must balance the necessity of glia in the short term with their harmful effects chronically.

## Acknowledgements

We thank G. Donahue for early guidance on analyzing RNAseq data; V. Lee and S. Narasimhan for helpful discussions on tau; R. Bonasio, L. Goodman, E. Lee, A. Perlegos, and K. Simeonov for manuscript feedback; F. Carranza and Z. Jin for technical assistance; M. Kayser, G. Ming, and D. Meany for helpful thesis feedback; S. Lindquist for continued inspiration; This work was supported by T32-AG000255, F31-NS111868 (to CNB) and R35-NS097275 (to NMB).

## Author contributions

Conceptualization, Investigation, Formal Analysis, and Visualization, C.N.B, under the Supervision of N.M.B; J.S. contributed to RNAseq study Conceptualization. Writing - Original Draft, C.N.B. Writing – Review & Editing, C.N.B and N.M.B; Funding Acquisition, C.N.B. and N.M.B

## Methods

### Fly work and lifespan analysis

All *Drosophila melanogaster* strains were maintained on standard cornmeal-molasses medium, flipped to fresh vials every 2 d and housed at 25°C on a 12 h light/dark cycle. For *geneSwitch* (inducible gal4-UAS)^79^ experiments, food was prepared with either 100 ul of RU486 (4 mg/ml in 100% EtOH; Sigma-Aldrich, M8046-1G) or vehicle (100% EtOH) pipetted onto food vials and allowed to dry for 24 h. For activating UAS-transgenes prior to injury, adult males were collected at eclosion and aged on RU486 or vehicle for 3 d, ensuring robust expression. For lifespans, flies were flipped to fresh food vials every 2 d and the number of dead/censored flies was recorded. See Supplementary Tables 1 and 2 for full experimental and genotype information.

### Localized head injury using dTBI model

Our *Drosophila* TBI (dTBI) paradigm was performed as described previously ^36^. Briefly, adult male flies were collected at eclosion and aged for 3 d. Flies were lightly anesthetized, collar restrained (n = 10 flies/collar) and allowed to fully recover from anesthesia. For dTBI injury, flies were individually subjected to closed-head injury at 35% (mild dTBI) or 45% (severe dTBI) head compression via controlled depression of a piezoelectric bar. For sham injury, sibling flies were handled in parallel (dTBI matched controls by vial) without head injury. An experimental cohort typically consisted of sibling sham or dTBI flies injured at the same time (n = 20 per vial). An independent experiment is defined as sham/dTBI experiments performed with non-sibling flies and with independent downstream processing. Flies were maintained on standard cornmeal-molasses medium, flipped to fresh vials every 2 d and housed at 25°C on a 12 h light/dark cycle.

### RNAseq experiment, library prep and sequencing

Severe and sham dTBI flies were injured and collected at 0, 1, 3, 6, 12, 24 hours and 3, 7, 10 and 15 days post-injury. Each timepoint had three independent experimental cohorts of n = 20 flies each. dTBI was performed between 09:00 - 12:00 to account for potential circadian effects. Whole flies were collected in 15 ml tubes, flash-frozen in liquid nitrogen then vortexed to separate heads from bodies. Total RNA from heads was extracted by TRIzol/Chloroform extraction (ThermoFisher, 15596026) with DNase treatment (ThermoFisher, AM1907). Total RNA integrity was checked by BioAnalyzer (no RIN score for flies). Dual-indexed libraries were generated using TruSeq Stranded mRNA Library Prep Kit with 96 Indices (Illumina, RS-122-2103) following the High Sample protocol. Libraries were prepared in two batches: samples from 0, 1, 3, 7, 10 and 15 days (Library 1) and samples from 0, 1, 3, 6, 12, 24 hours (Library 2). Library preparation failed for one 15 dpi dTBI replicate so this sample, along with the matched sham, were excluded from further analyses. To aid bioinformatic correction of batch effects introduced during library prep, libraries were prepped from 0 and 1 d samples twice (Library 1 and 2), using the same total RNA. Library concentration was measured by Qubit and quality checked for insert size and primer-dimer contamination (200-300 bp; TapeStation). Libraries were pooled and sequenced on a NextSeq, using the NextSeq 500/550 High Output Kit v2 with 75 cycles (Illumina, FC4042005).

### RNAseq alignment and differential expression analysis

Demultiplexed reads passing QC filter (Q > 30) were obtained from BaseSpace then merged across sequencing lanes for each sample, with approximately 13-20 million reads total per sample. Single-end reads were aligned to the fly genome using HISAT2^80^. The HISAT2 index was built from FlyBase’s *Drosophila melanogaster* reference genome r6.17. BAM files for each sample were merged across sequencing runs (Picard). Reads that uniquely aligned to exonic regions were counted with HTSeq^81^ with the union setting to produce a count matrix for differential expression analysis using the DESeq2^82^ package in the R environment.

Principal component analysis was used to determine batch effects due to library preparation as follows. First, variance stabilized count data for all samples was visualized with plotPCA(). When samples were colored by library preparation, PC1 was associated with library preparation. Removing library effects with limmaRemoveBatchEffect() restored PC1 and PC2 to injury status and/or time post-injury. Thus, library was included in the DESeq2 design. Libraries prepared twice (0, 1d) did not cluster by condition after batch effect correction thus were treated as biological, not technical, replicates. To identify differentially expressed genes at each time post-injury, samples were reclassified into groups based on injury condition and post-injury time (i.e: TBI_15dpi, TBI_1dpi). The design model formula was “∼library + group”. Pairwise comparisons were made between sham and dTBI samples at each time using “contrast=c(“group”), with an alpha cutoff of 0.05 with lfcShrink() applied.

### Motif and pathway enrichment analyses

*De novo* motif enrichment analysis was performed with HOMER’s^83^ findMotifs.pl using the provided fly promoter set. All default parameters were used with the following exceptions: promoter region (−1000, 300 bp from TSS) and background promoter frequency. Background promoter frequency was derived from our RNAseq data and consisted of all expressed genes (non-zero read count; n = 16,201). Enrichment for *de novo* motifs was performed on significantly up or down-regulated genes at each time post-injury. Motifs passing a false positive cutoff (p < 1.0e-10) were matched to the best-known *Drosophila* motif or motif binding factor. Motif target genes were obtained using HOMER’s annotatePeaks.pl. Enrichment for KEGG and Reactome pathways was determined using the interactive online tool FlyMine ^84^. Curated gene sets were analyzed for pathway enrichment with Benjamini-Hochberg correction, using the same RNAseq defined background gene set.

### Real-time quantitative PCR (RT-qPCR)

Total RNA was isolated from dissected brains (n = 9 brains per experimental cohort) by RNeasy Mini Kit (Qiagen, 74104), with on-column removal of genomic DNA (Qiagen, 79254). Complementary DNA was prepared from total RNA (Applied Biosystems, 4368814) then quantified by Qubit ssDNA Assay (Invitrogen, Q10212). qPCR reactions were set up using Fast SYBR Green reagents (ThermoFisher, 4385612), 384-well plates, 20 ng of cDNA per reaction, and analyzed on a ViiA 7 Real-Time PCR System (Applied Biosystems). Relative expression was determined using the ΔΔCT method. For each sample, mean CT values were determined from 2-3 technical replicates. ΔCT was determined relative to the housekeeping gene, *RpL32*, which was unaffected by dTBI according to RNAseq data. ΔΔCT was then calculated as fold-change relative to the control group, which is noted in the Fig. legend. qPCR primers were sourced from FlyPrimerBank^85^ or prior publications, BLAST’d against the fly (and human) genome for specificity and optimized by serial dilution curve and melt curve analysis. See Supplementary Table 3 for primer information.

### Whole mount brain immunofluorescence

Fly brains were dissected in cold phosphate buffered saline (PBS), as described^86^. Brains were fixed in 4% paraformaldehyde in PBS for 55m at room temperature (RT) then permeabilized overnight in 0.5% Triton-X in PBS (PBST) at 4°C. Genotypes with *TREdsRed, MREdsRed*, or *UAS-mCD8-GFP* were imaged without further antibody staining. Brains were blocked with 5% normal goat serum in PBST for 1h at RT then incubated with 1° antibody overnight at 4°C (see Supplementary Table 3 for antibody information). Brains were washed in PBST then incubated with fluorescently conjugated 2° antibody for 1h at RT. See Table S4 for antibody information. Brains were counterstained with Hoechst (0.10 mg/ml in PBS) for 15m, cleared in mounting media (20 mM Tris pH 8.0, 0.5% N-propyl gallate, 80% glycerol, PBS), mounted and cover slipped. Brains were imaged by confocal microscopy (Leica DM 6000 CS) with identical laser power and gain settings across experiments. For whole brain imaging, images were acquired throughout the full brain at 2 *μ*M steps, 1024×1024 resolution by 10X, 20X (dry) or 60X (oil) objectives. For dsRed quantification, dsRed was measured in FIJI as raw integrated density in scaled images of the z-stacked brain. For colocalization of dsRed with repo and elav, nucleated dsRed+ cells were counted using the cell counter feature. Colocalization with repo or elav was determined. Image acquisition and analysis was performed blind to sample identity.

### Western immunoblot

Brains were rapidly dissected in cold PBS (n = 8 per biological replicate), then homogenized in 5 *μ*l sample buffer per brain (1x Laemmli Buffer [Bio-Rad, 161-0737], 1x cOmplete mini EDTA-free protease inhibitor cocktail, 1 mM PMSF [Sing, P7626], 50 *μ*l BME [Sigma, M6250]). Samples were denatured (98°C for 3 m) prior to loading onto 4-12% Bis-Tris gel. Volume equivalent of 1 brain per sample was run in 1x MES buffer, transferred to 0.45 *μ*M nitrocellulose membrane overnight by electrophoresis. Membranes were stained by ponceau stain to confirm transfer. Membranes were blocked in 3% bovine serum albumin in 1X Tris-Buffered Saline, 0.1% Tween® 20 Detergent (TBST), incubated in primary antibody overnight at 4°C. See Table S4 for antibody information. Blots were incubated with 1:5000 dilution of species-appropriate HRP-conjugated secondary antibody for 1 h at RT, then detected by ECL (Cytiva (Formerly GE Healthcare Life Sciences), RPN2232) using an Amersham Imager 600. Quantification was performed in FIJI by ROI. Sample protein was normalized to a loading control, alpha Tubulin.

### Paraffin embedding of fly heads and brain vacuole quantification

Heads were rapidly decapitated under anesthesia then fixed in Bouin’s solution (Sigma-Aldrich, HT10132) for 5 days. Fixation was terminated by submersion in leaching buffer (50mM Tris pH 8.0, 150mM NaCl) for 1 h at RT. Heads were then processed through graded EtOH dehydration at the following times and concentrations: 30 m 70%, 30 m 90%, 30 m 95%, 30m 100%, 30 m 100%, followed by xylenes (2x, 30 m) and finally fixed in paraffin. Heads were blocked and sectioned into 8 uM thick ribbons. Ribbons were deparaffinized by heating at 65°C for 1 h followed by washes in histoclear I and II (VWR; 101412-876, 10141-8822) for 5 m. For assessing brain vacuolization, sections were mounted with Cytoseal XYL (ThermoFisher, 8312-4). Sections were imaged on a Leica DFC360 FX under 10x objective, 1.2x magnification with fixed exposure settings. Fly brain tissue is auto-fluorescent for pigmented eyes thus signal was detected using an I3 filter cube. Brain vacuolization was quantified in FIJI as follows: for each brain, vacuolized area and total brain area were averaged across 5 non-consecutive sections (every 3^rd^ section) to determine % of brain vacuolization. Vacuole area was set by manual thresholding. Image acquisition and analysis was performed blind to sample identity.

### Immunofluorescence on paraffin sections

Sections were rehydrated to water as follows: 100% EtOH for 3 m, 100% EtOH for 3 m, 95% EtOH for 1 m, 80% EtOH for 1 m, H_2_O for 5 m. Antigen retrieval was performed by boiling slides in citric acid buffer (Vector; H-3300) at 95°C for 40 m. Sections were then permeabilized in PBS with 0.1% Tween-20 (PBSTw), blocked in 3% bovine serum albumin (BSA) for 1 h at RT then incubated in 1° antibody overnight at 4°C. Sections were washed (2x, PBSTw) then incubated in 2° antibody for 1 h at RT. See Table S4 for antibody information. Sections were washed in PBSTw, rinsed in deionized water then coverslipped with mounting media. For phospho-tau puncta quantification, sections were imaged on a Leica DFC360 FX under a 10x objective, 1.2x magnification and identical exposure settings. Puncta were counted in FIJI using the cell counter. Puncta were counted in one brain hemisphere across 5 non-consecutive sections (every 3^rd^ section) to calculate puncta per 100^2^ *μ*M^2^. High resolution representative images were acquired using a 20x and 60x objective (oil immersion) on a Leica DM6000 CS. Image acquisition and analysis was performed blind to sample identity.

### Phosphatase shift assay with SDD-AGE

Whole flies (n = 30 per condition) were flash frozen then vortexed to collect heads. Tissue was homogenized in 100 *μ*l cold RIPA buffer (Pierce, 8990) supplemented with cOmplete mini EDTA-free protease inhibitor cocktail (Roche, 11836170001). Protein abundance was measured by Qubit. Samples were diluted to 4 *μ*g/*μ*l then split into two 50 *μ*g samples (treated and untreated). Phosphatase treated samples were enzymatically treated for 3 n at 37°C shaking (1.25 *μ*l 10 *μ*M MnCl_2_, 1.25 *μ*l 20X NEB buffer, 2.5 *μ*l lambda phosphatase enzyme). Samples were run on 1.5% agarose gel with 0.1% SDS at 100 mV. Protein was transferred to 0.45 *μ*M nitrocellulose membrane overnight^87^. Membrane was then processed by immunoblotting as above. See Supplementary Table 4 for antibody information.

### Human TBI analysis

Pathology data for brain samples from male donors (n = 191 samples, n = 59 unique donors) was obtained from the Allen Institute for Brain Science^88,89^. Corresponding RNAseq data was obtained as aligned BAM files. Exonic read counts were calculated using the R package Rsubread (GTF file was GRCh38.p2). Differential gene expression analysis (FDR < 0.10) was performed in DESeq2 as above, with the following modifications. Model design (injury + brain region) was determine after principal component analysis. PCA revealed extreme outliers without an obvious pattern (n = 37; see **Fig. S7B**). Removal resulted in all samples clustering by brain region along PC1 and PC2, allowing for inclusion of samples from distinct brain regions from the same donor (n = 161; 80 non-TBI and 81 TBI samples). Motif enrichment analysis against known motifs was performed using HOMER as above. Enriched molecular signature data base (mSigdB) gene sets and gene ontology terms were obtained from HOMER’s hypergeometric comparison (p < 1.0e-7).

### Statistical analyses and data visualization

All statistical analyses and data visualization were performed in the bash (RNAseq alignment and HOMER analysis) or R environment using RStudio and the following packages: annotables, biomaRt, cowplot, DESeq2, ggplot2, ggrepel, ggridges, ggsignif, ggpubr, homerkit, lawstat, RColorBrewer, RBase, survminer, tidyverse. Specific tests are indicated in the Fig. legends. Data were analyzed by either parametric or non-parametric tests based on the sample size, homogeneity of variance (Levene’s test) and normality (Shapiro-Wilks test).

### Data Availability

The differential gene expression data generated in this study are included in this published article (and its supplementary information files). Sequencing data that support the findings of this study have been deposited in Gene Expression Omnibus (accession number to be added before publication). Additional data that support the findings of this study are available from the corresponding author upon reasonable request.

---

**Extended Data Fig 1.**
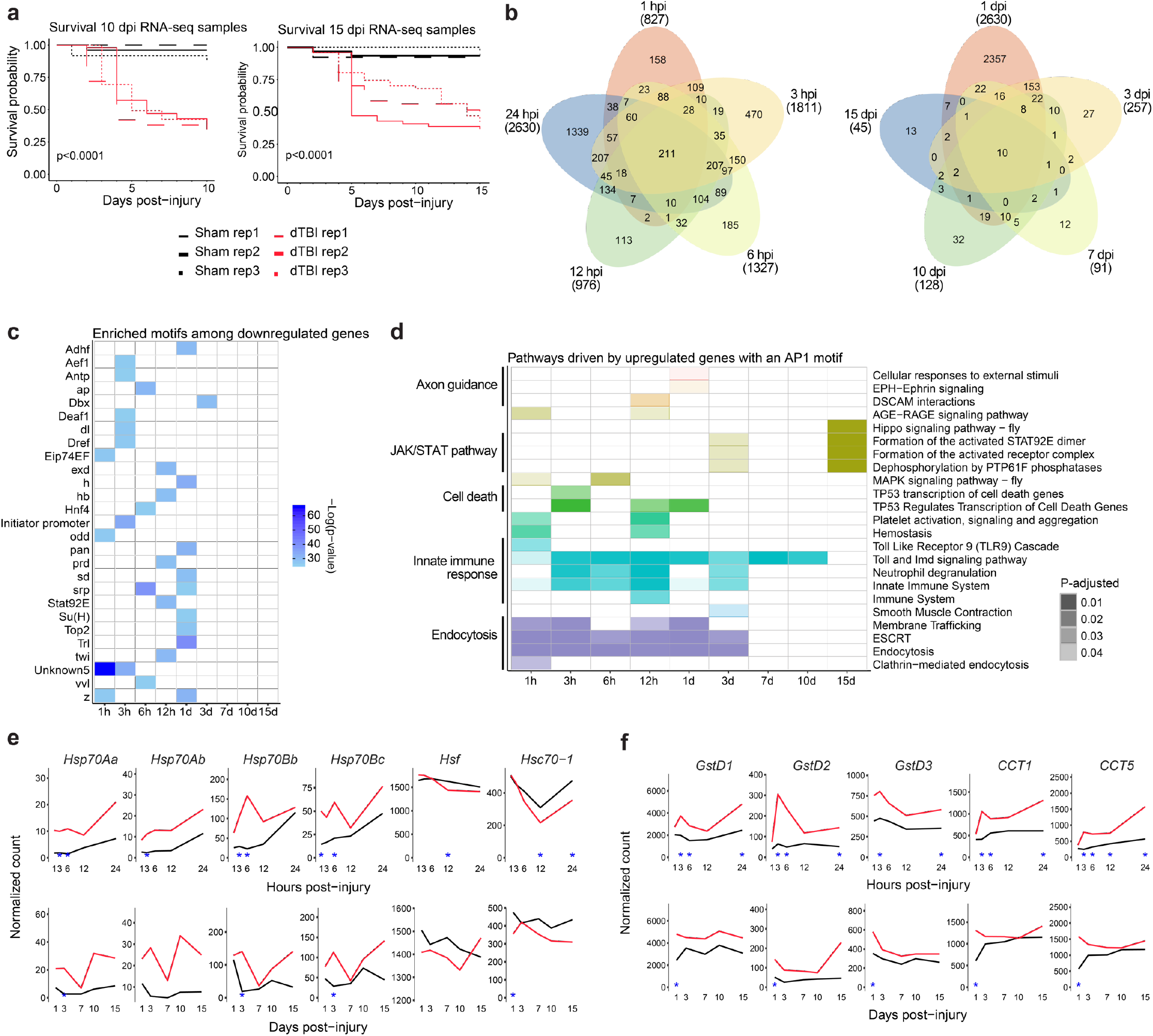
A lasting AP1 transcriptional response to TBI. **a**, Kaplan-Meier survival curve for 10 and 15 dpi RNAseq replicates. **b**, Venn diagram showing the number of common and distinct differentially expressed genes (FDR<0.05) between post-injury times. Value in parenthesis shows total (up and down) number of DE genes for a given timepoint. **c**, Results for HOMER *de novo* motif enrichment among downregulated genes (FDR<0.05). **d**, Tile plot showing Reactome pathways enriched among upregulated genes with a predicted AP1 motif (FDR<0.05). Presence of a colored tile indicates enrichment at a given post-injury time. Tile opacity encodes significance. Color corresponds to the parent process, as defined by the Reactome annotation database (annotated on left). **e**, Average sham (black) and severe dTBI (red) expression of genes related to the heat shock response at ≤1 dpi (top) and ≥1 dpi (bottom; blue star indicates FDR < 0.05 at a given time). **f**, Average sham (black) and severe dTBI (red) expression of canonical stress response genes at ≤1 dpi (top) and ≥1 dpi (bottom; blue star indicates FDR < 0.05 at a given time). See Supplementary Table 1 for genotypes.

**Extended Data Fig. 2.**
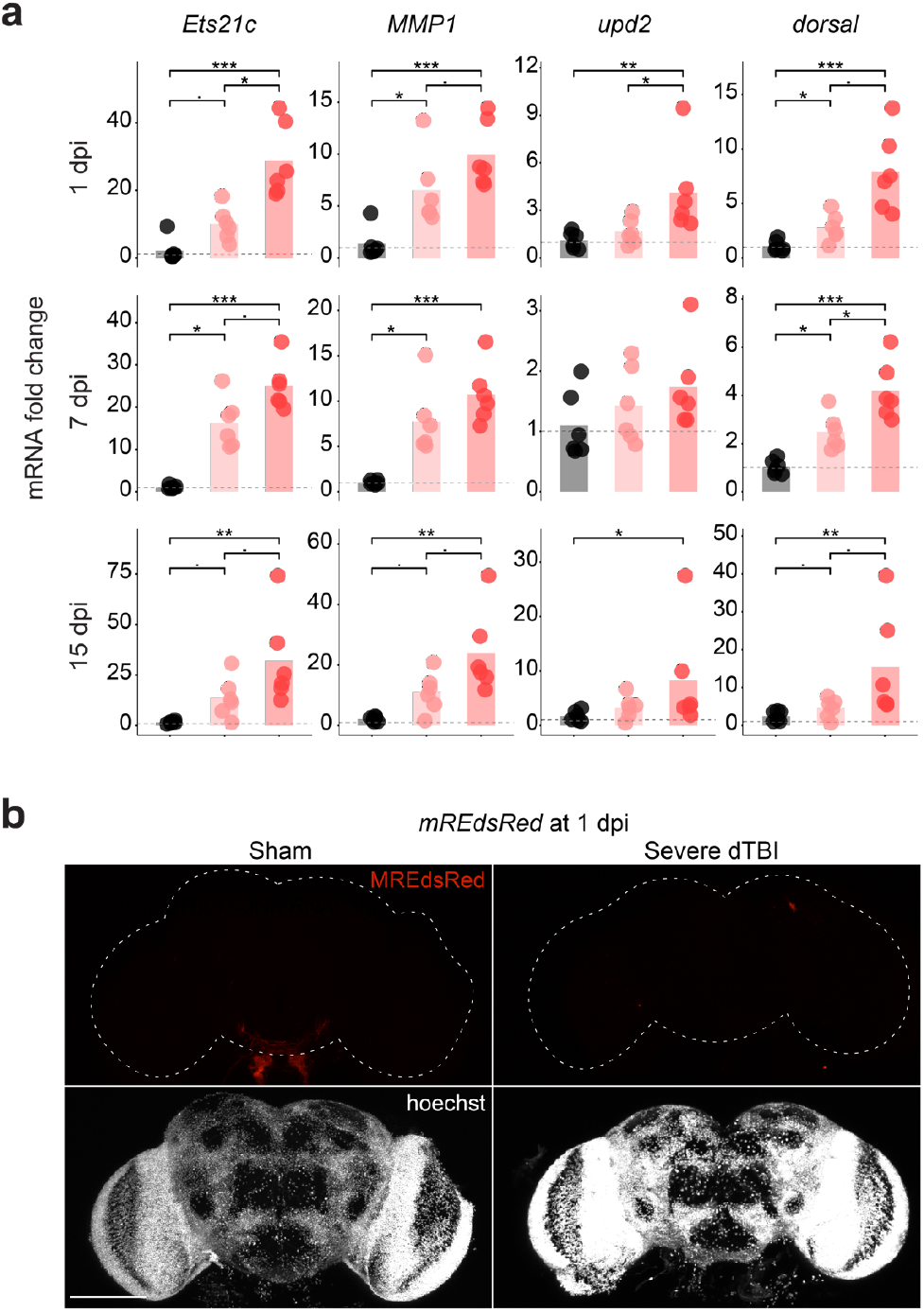
Sustained and severity-dependent activation of AP1. **a**, Mean relative expression of select RNAseq predicted AP1 genes by RT-qPCR at 1, 7 and 15 dpi across injury conditions (each point = 1 biological replicate, 9 dissected brains; n = 6 biological replicates per condition; Kruskal–Wallis test with Dunn’s multiple comparison test and Holm adjustment for each gene). **b**, Representative z-stacked whole mount brains at 1 dpi in flies with dsRed expressed under a mutated TRE-promoter (*MREdsRed*; representative of n = 9 - 11 brains per condition from two independent experiments). Statistical annotations are ****p<0.0001, ***p<0.001, **p<0.01, *p<0.05. All scale bars 100 #m. See Supplementary Table 1 for genotypes.

**Extended Data Fig. 3.**
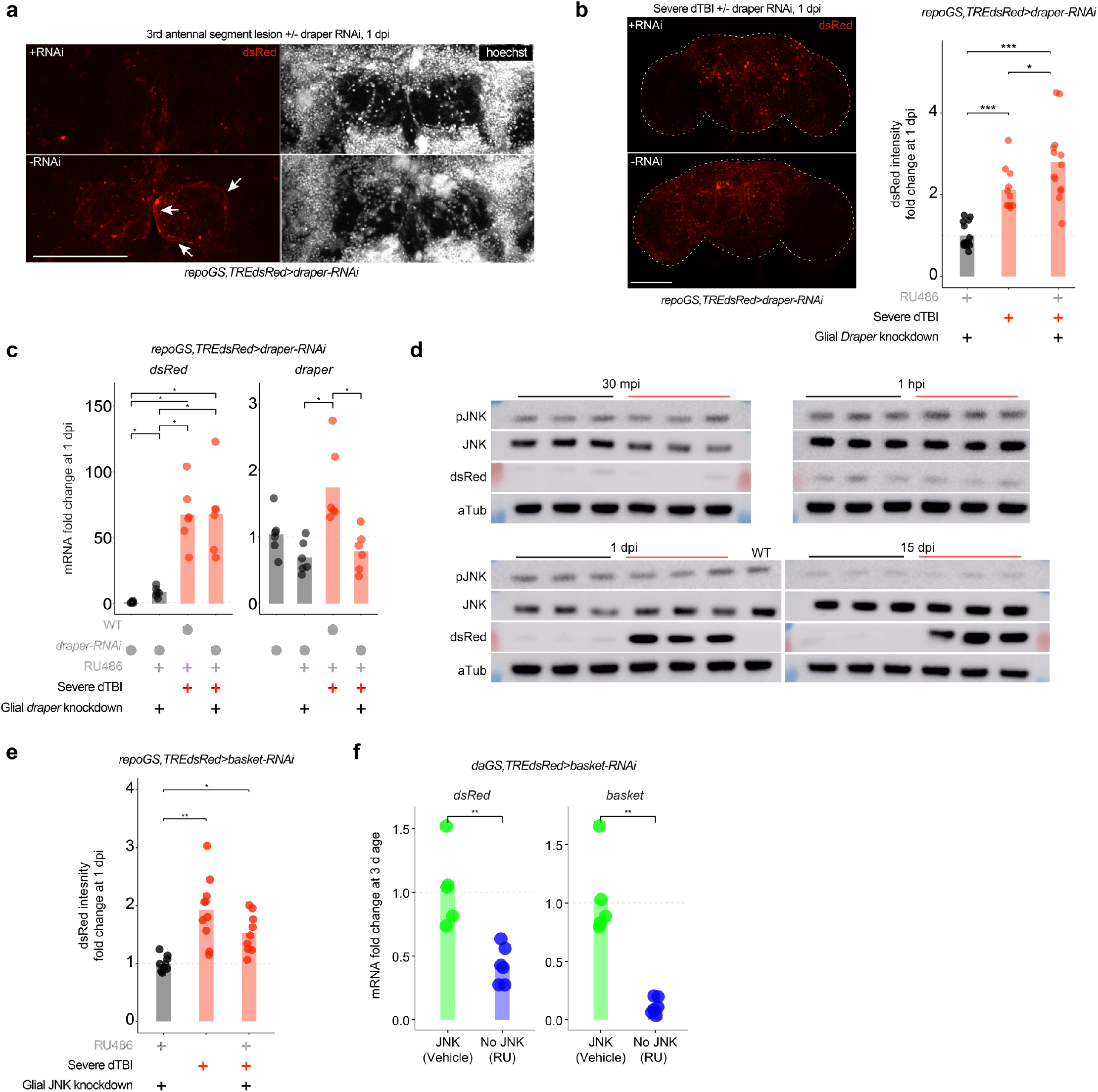
Glial AP1 activation is draper and JNK-independent. **a**, Representative z-stack of the first 6 slices (2 #m each) of the antennal lobe in flies with (RU; top) and without (vehicle; bottom) *draper-RNAi* in glia. White arrows highlight hypertrophic glial processes. Image shown is 1 day after 3^rd^ antennal segment ablation (abbreviated AL injury; images are representative of n = 7 - 9 brains per condition). **b**, Left, representative z-stacked whole mount brains at 1 day post-dTBI with (RU; top) and without (vehicle; bottom) *draper-RNAi* in glia. Right, quantification of dsRed immunofluorescence fold change, relative to left most condition (each point = 1 brain; n = 11 - 14 brains per condition pooled from two independent experiments; p = 2.17e-06, Kruskal–Wallis test with Wilcoxon rank sum test). **c**, Mean relative expression of *dsRed* and *draper* mRNA by RT-qPCR at 1 dpi, with or without glial *draper-RNAi* (each point = 1 biological replicate, 9 dissected brains; n = 6 biological replicates per condition; Kruskal–Wallis test with Dunn’s multiple comparison test and Holm adjustment). **d**, Western immunoblots for all replicates of Fig. 4a. Samples are biological replicates. WT is *5905*. **e**, Quantification of dsRed immunofluorescence fold change for representative whole mount brains shown in Fig. 4b, relative to left most condition (each point = 1 brain; n = 9-10 brains per condition pooled from two independent experiments; p = 0.000238, Kruskal–Wallis test with Wilcoxon rank sum test). **f**, Mean relative expression of *dsRed* and *basket* mRNA by RT-qPCR in 3 d old whole flies, with or without JNK RNAi expressed under a ubiquitous driver, *daughterlessGeneSwitch* (each point = 1 biological replicate, 20 whole flies; n = 6 biological replicates per condition; Student’s t-test for each gene). Statistical annotations are ****p<0.0001, ***p<0.001, **p<0.01, *p<0.05. All scale bars 100#m. See Supplementary Table 1 for genotypes.

**Extended Data Fig. 4.**
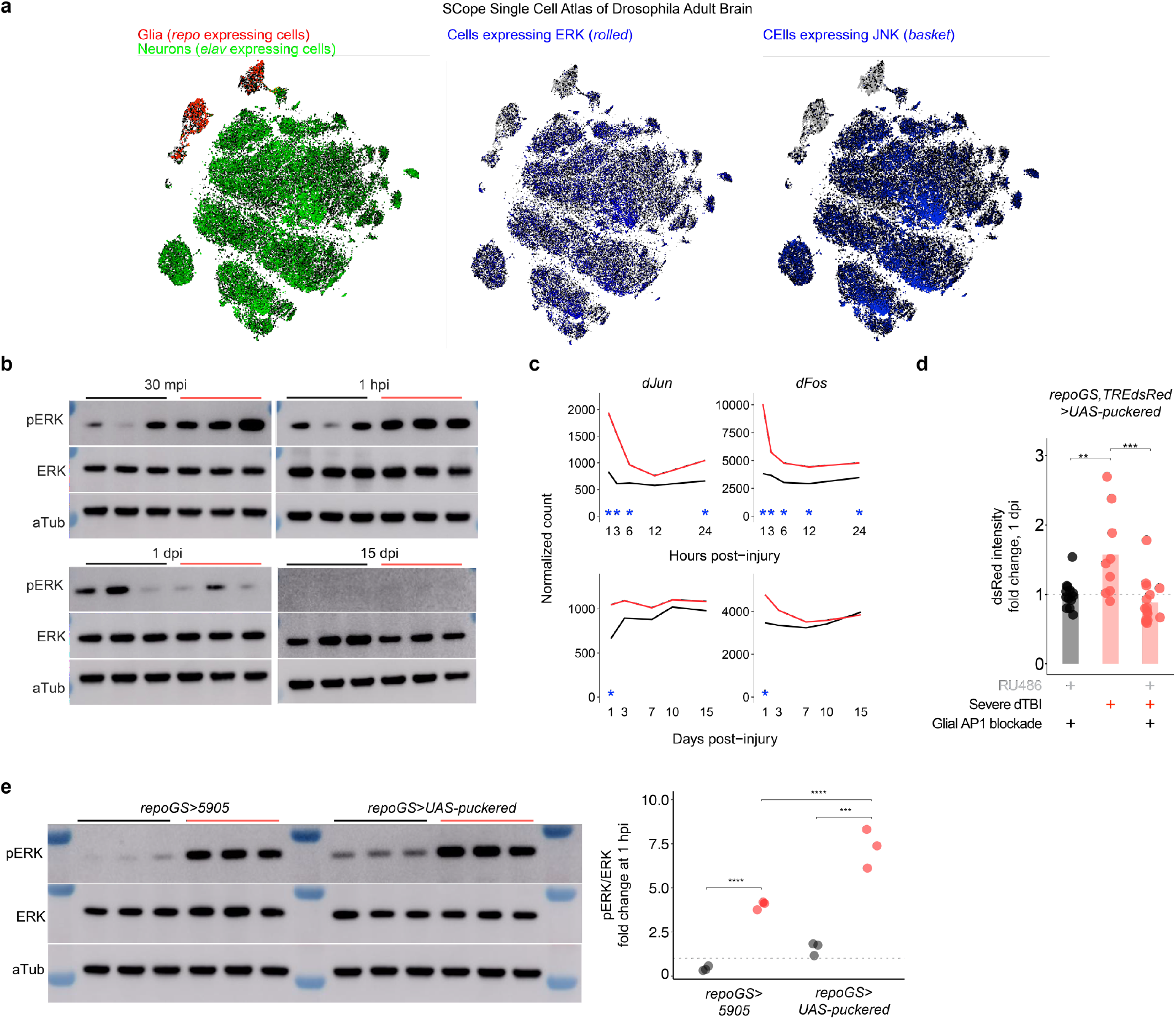
AP1 activation requires ERK. **a**, Images from Scope Single Cell Atlas of the *Drosophila* adult brain showing relative expression levels of *repo, elav, rolled* (ERK), and *basket* (JNK). ERK is detected in a greater proportion of repo+ cells than JNK. **b**, Western immunoblots for all replicates of Fig. 4c. Samples are biological replicates. Genotype is *TREdsRed*. **c**, RNAseq data showing average sham (black) and severe dTBI (red) expression of *dFos* and *dJun* at ≤1 dpi (top) and ≥1 dpi (bottom; blue star indicates FDR < 0.05 at a given time). **d**, Quantification of dsRed immunofluorescence fold change for representative whole mount brains shown in Fig. 4g, relative to left most condition (each point = 1 brain; n = 9-15 brains per condition pooled from two independent experiments; p = 0.0040, Kruskal–Wallis test with Wilcoxon rank sum test). **e**, Glial expression of the MAPK phosphatase, *puckered*, does not reduce pERK levels at 1 hpi. Left, full western blots. Right, quantification of pERK/ERK ratio by western immunoblot (each point = 1 biological replicate, 8 dissected brains; n = 3 biological replicates per condition/genotype; p = 0.0159, two-way ANOVA). Statistical annotations are ****p<0.0001, ***p<0.001, **p<0.01, *p<0.05. See Supplementary Table 1 for genotypes.

**Extended Data Fig. 5.**
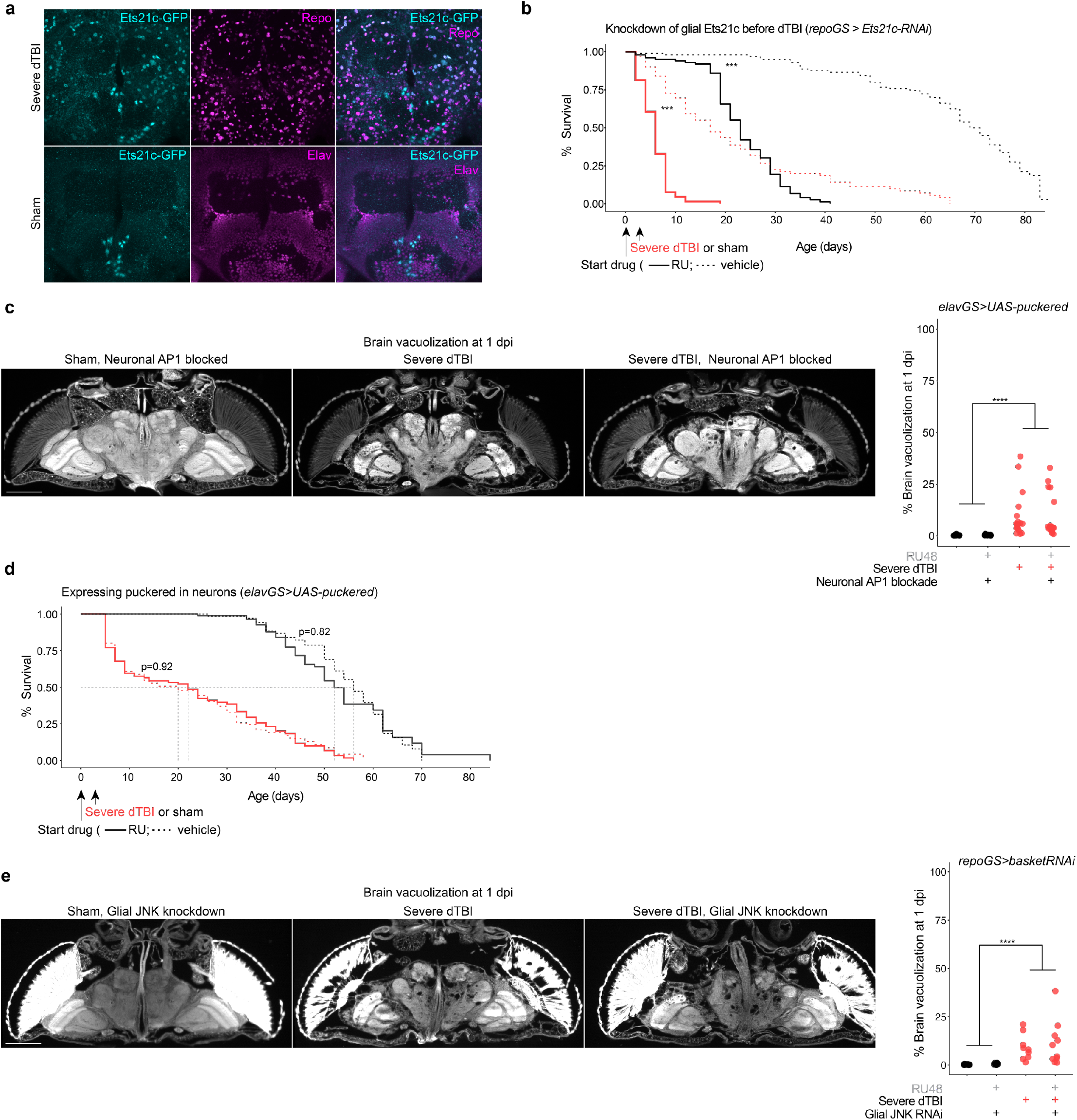
Glial AP1 is required for TBI recovery. **a**, Representative z-stacked whole mount brains focused on the central brain at 1 dpi in flies expressing a GFP-tagged variant of the AP1 target gene, *Ets21c*. Ets21c-GFP is detected in a handful of neurons at baseline but is dramatically upregulated in glia by 1 dpi (representative of n = 11-15 brains per condition, two independent experiments). **b**, Post-injury survival with (RU; dashed line) or without (vehicle; solid line) *Ets21c-RNAi* expressed in glia (n = 100 per condition, 5 vials of 20 flies; p < 0.0001, Kaplan-Meier analysis with log-rank comparison). **c**, Representative brain vacuolization at 1 dpi under sham and dTBI conditions, with (RU) or without (vehicle) JNK RNAi expression in glia. Quantification on right, expressed as % of total brain area that is vacuolized (each point = 1 brain; n = 9-10 brains per condition; two-way ANOVA revealed a non-significant interaction but a significant effect of dTBI, p = 6.1e-05). **d**, Representative brain vacuolization at 1 dpi under sham and dTBI conditions, with (RU) or without (vehicle) *puckered* expression in neurons. Quantification on right, expressed as % of total brain area that is vacuolized (each point = 1 brain; n = 18-19 brains per condition pooled from two independent experiments; two-way ANOVA revealed a non-significant interaction but a significant effect of dTBI, p = 3.55e-06).**e**, Post-injury survival with (RU; dashed line) or without (vehicle; solid line) *puckered* expression in neurons (n = 100 per condition, 5 vials of 20 flies; p < 0.0001, Kaplan-Meier analysis with log-rank comparison). Statistical annotations are ****p<0.0001, ***p<0.001, **p<0.01, *p<0.05. All scale bars 100#m. See Supplementary Table 1 for genotypes.

**Extended Data Fig. 6.**
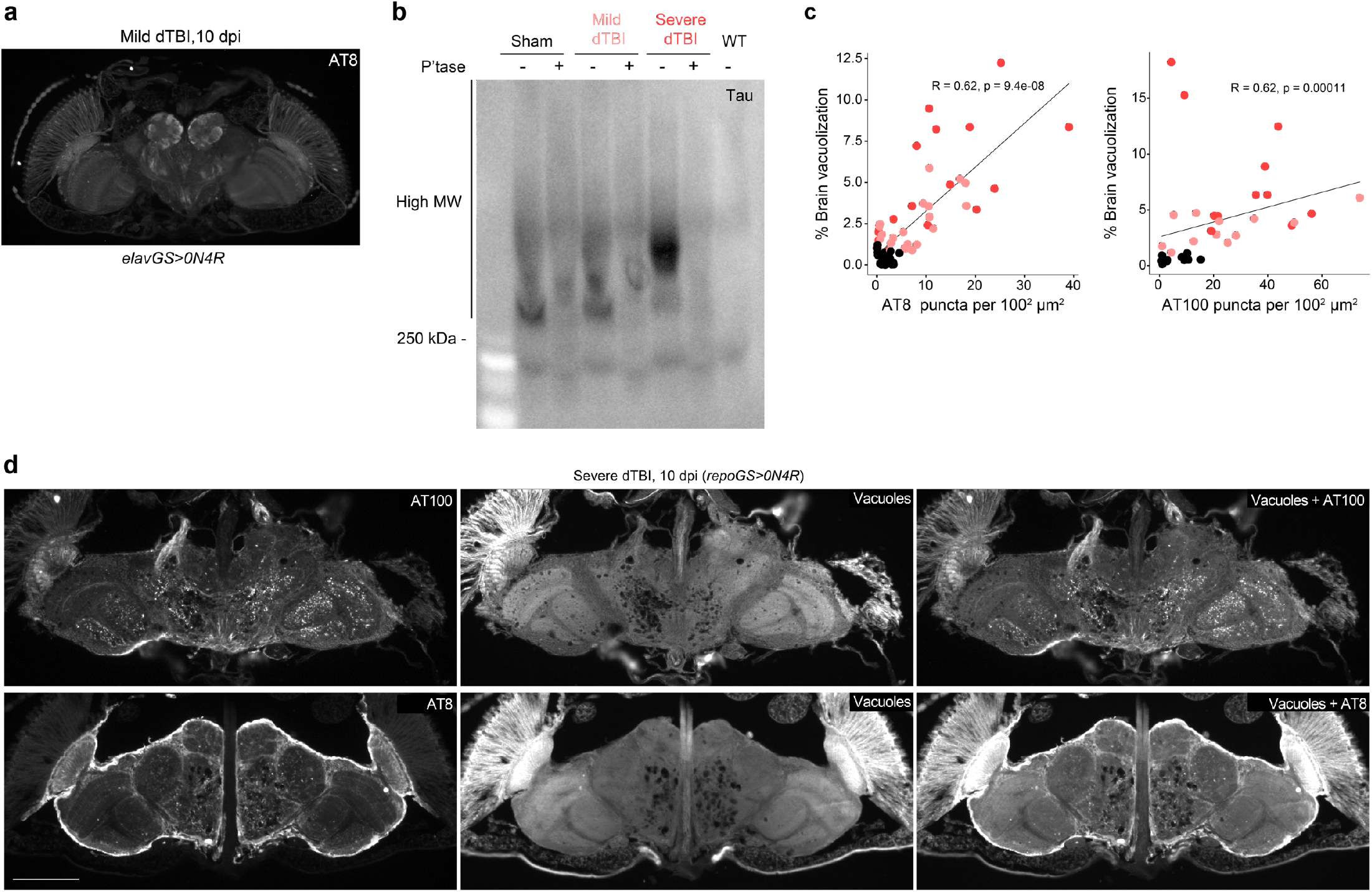
Sustained glial AP1 activity drives human tau pathology. **a**, Representative section of paraffin-embedded head at 10 dpi stained for AT8 when human tau is expressed in neurons (representative of n = 7 animals). **b**, Membrane immunostained for human tau in phosphatase treated (+) and untreated (-) protein samples isolated from the heads of 5 dpi (*repoGS>0N4R*) or WT (*5905*) flies (n = 30 per sample). **c**, Spearman correlation between percentage of total brain area vacuolized and number of AT8+ puncta (top) or AT100+ puncta (bottom) at 5 and 10 dpi (combined) in sham, mild or severe dTBI. Data shown is an alternative representation of Fig. 6a. **d**, Representative sections of paraffin embedded heads immunostained for AT100 (top row) or AT8 (bottom row), highlighting the concentration of phosphorylated tau puncta in neuropil and around vacuoles. Vacuoles are visualized by autofluorescence. Statistical annotations are ****p<0.0001, ***p<0.001, **p<0.01, *p<0.05. All scale bars 100#m. See Supplementary Table 1 for genotypes.

**Extended Data Fig. 7.**
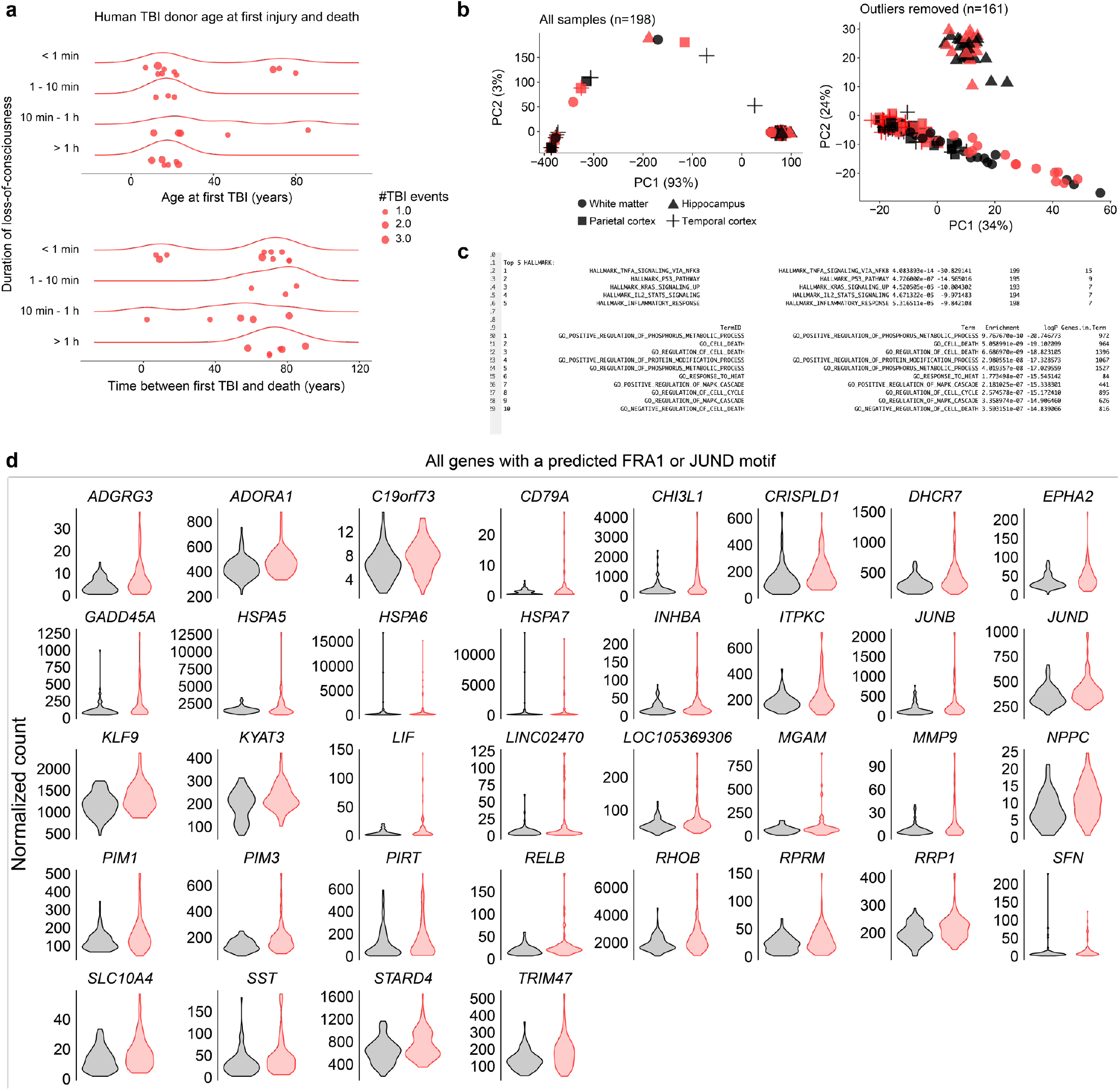
Evidence of AP1 activity decades after moderate human TBI. **a**, TBI exposure history for donors; the same data is visualized two ways. Top shows age at first TBI, with most injuries occurring before age 30. Donors are further separated by duration of loss-of-consciousness. Total lifetime TBI events for each donor are shown by point size. Bottom shows the duration of time between first TBI and death. **b**, PCA of all samples (left) compared PCA of samples with outliers removed (right). Each point represents 1 tissue sample. Shape encodes brain region. **c**, Top 5 Molecular Signature Data Base (MSigDb) Hallmark gene sets and top 10 gene ontology (GO) terms enriched among upregulated genes (FDR<0.10) in TBI donors. **d**, Violin plots showing expression of all predicted FRA1/JUND genes in non-TBI (black; n samples = 81) and TBI (red; n samples = 80) exposed donors (FDR < 0.10).

